# A clinical grade neurostimulation implant for hierarchical control of physiological activity

**DOI:** 10.1101/2025.10.21.683630

**Authors:** Moaad Benjaber, Mayela Zamora, Robert Toth, John E Fleming, Kei Landin, Victoria S Marks, Rachel A Crockett, Jae-Wook Ryou, Alceste Deli, Alexander L Green, Rory J Piper, Martin M Tisdall, Derk-Jan Dijk, Nicholas D Schiff, Andrew Sharott, Keith P Purpura, Jonathan L Baker, Joram J van Rheede, Timothy J Denison

**Affiliations:** Institute of Biomedical Engineering, Department of Engineering Science, University of Oxford, Oxford, UK; Medical Research Council Brain Network Dynamics Unit, Nuffield Department of Clinical Neurosciences, University of Oxford, Oxford, UK; Feil Family Brain and Mind Research Institute, Weill Cornell Medicine, New York, USA; Academic Neurosurgery Unit, Neuroscience and Cell Biology Research Institute, City St. George’s University of London, London, UK; Nuffield Department of Surgical Sciences, University of Oxford, Oxford, UK; Department of Neurosurgery, Great Ormond Street Hospital, London, UK; UCL Great Ormond Street Institute of Child Health, University College London, London, UK; Surrey Sleep Research Centre, University of Surrey, Guildford, UK; UK Dementia Research Institute, Care Research and Technology Centre at Imperial College London and the University of Surrey, Guildford, UK

## Abstract

Bioelectronic implants for neurostimulation aim to steer disordered neurophysiological processes back towards a healthy state. However, physiology is subject to biological rhythms, including the circadian rhythm and the sleep-wake cycle. These predictable rhythms affect disease symptomatology, biomarkers used in closed-loop therapies, and a physiological system’s expected response to stimulation. Therefore, therapeutic devices should incorporate feedforward elements to align algorithm parameters with predictable changes in physiological state, as a parallel of physiological rheostatic control. Here we introduce the DyNeuMo-2c, the first clinical-grade implant capable of delivering closed-loop neurostimulation while flexibly changing its functional configuration according to time of day. The device can chronically measure brain activity and motion state to track potential biomarker patterns in natural, out-of-clinic settings, allowing identification and targeting of patient-specific chronotypes. The system implements a hierarchical control flow, with baseline therapy set by a circadian scheduler, and adaptive policies layered to take effect based on specific biomarkers indicating patient and disease state. Using a benchtop validation setup, we demonstrate that the system has the required capabilities for delivering time-contingent closed-loop therapy in two established clinical use cases: Parkinson’s disease and epilepsy. Next, we deploy the system *in vivo* to deliver closed-loop deep brain stimulation in a healthy non-human primate model of vigilance, highlighting the importance of synchronisation between device operation and physiological state in various conditions (task performance, unconstrained behaviour, and sleep). Time-of-day-dependent adaptation of closed-loop stimulation enabled modulation of both vigilance and behaviour. Overall, the novel device architecture provides a proof-of-concept for delivering time-contingent therapy in chronic therapeutic settings where biological rhythms are of key importance.

## 1. Introduction

Deep brain stimulation (DBS) is a bioelectronic medical therapy in which an implanted system delivers electrical stimulation to deep brain structures to reduce pathological brain activity. This approach is proving fruitful in an increasing number of neurological and psychiatric conditions^1,2^. While most DBS systems continuously apply the same stimulation parameters (**Figure 1a**), recent technological developments have enabled the delivery of closed-loop therapy, in which stimulation is adjusted according to a feedback variable (biomarker) correlated with symptoms (**Figure 1b**)^3–6^. Such closed-loop capabilities push neuromodulation therapy into the realm of control theory^7,8^. According to the good regulator theorem, ‘*every good regulator of a system must be a model of that system’*^9^. In the context of DBS, this means we need implantable devices that can anticipate their interaction with neural dynamics at multiple clinically relevant timescales^10^.

**Figure 1:**
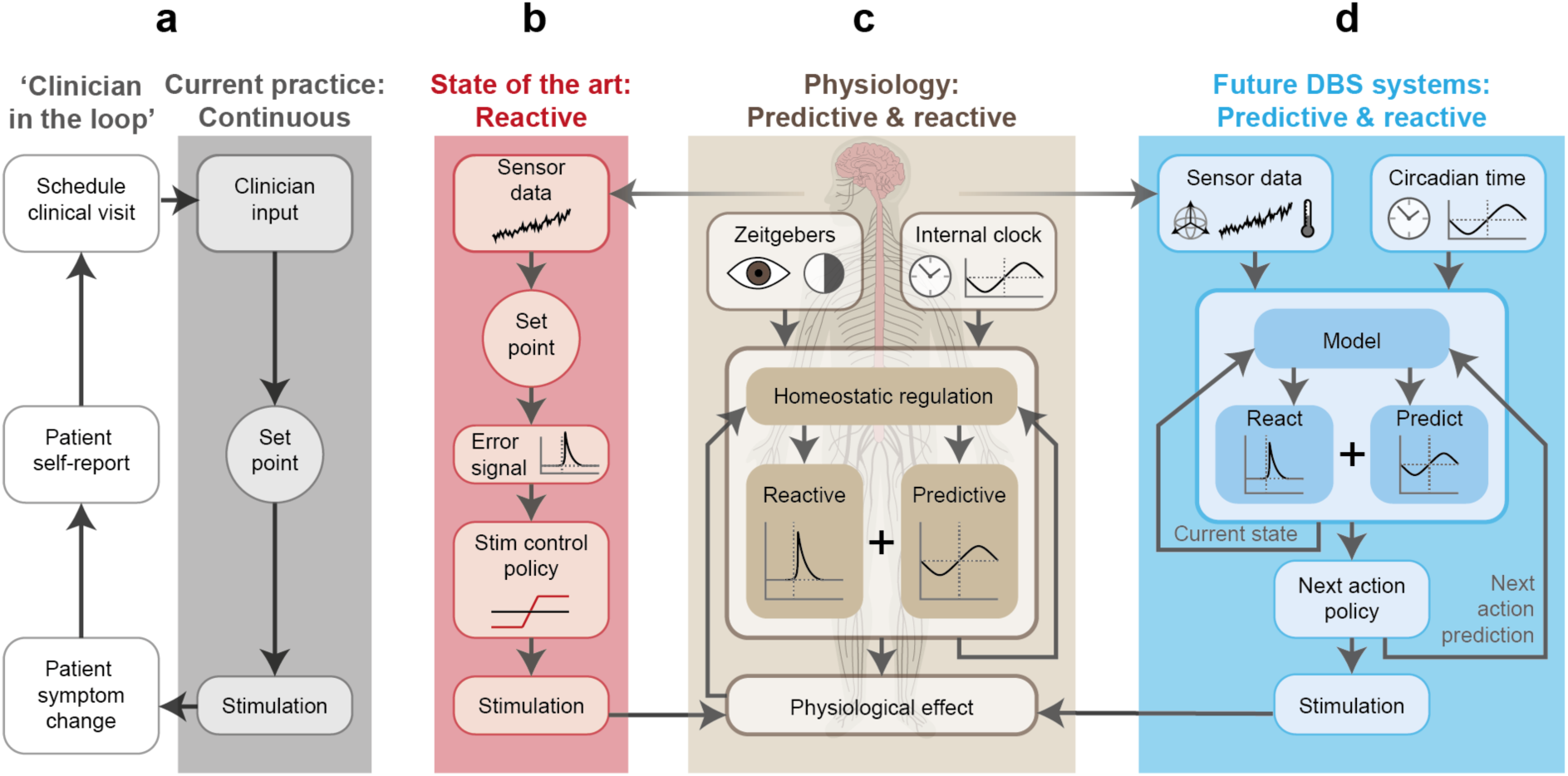
Introducing rheostatic principles for the control of medical devices. **a)** In standard DBS practice, the clinical team sets stimulation parameters during a daytime programming session with the patient, seeking to balance symptom control and side effects, as well as their modulators such as medication regime. The therapy then runs with these parameters continuously, 24 hours per day. While parameters can be adjusted based on patient feedback, this requires scheduling a new clinical appointment for DBS reprogramming. **b)** Emerging automated algorithms such as adaptive DBS record neural biomarker of therapeutic need and automatically adjust stimulation dose attempting to keep the marker within a specified range. **c)** Physiology incorporates both reactive (feedback) loops and predictive (feedforward) elements such as the circadian system that aim to prepare our physiology for a future anticipated state of the world and the body. Human nervous system adapted from original by Zane Mitrevica (https://doi.org/10.5281/zenodo.4646639). **d)** Future DBS therapies could take inspiration from our body’s rheostatic regulatory processes in which setpoints vary by incorporating both reactive and time-dependent predictive control elements. Sensor data can inform an internal model about the current state of the patient, while past sensor data, time of day and circadian phase can be used to predict the most likely future state and therefore anticipate therapeutic need.

While physiological homeostasis is commonly described in terms of feedback control – reacting to disturbances in the body’s internal environment – there is also a substantial feedforward element to physiological regulation. Anticipatory adjustments to physiological setpoints based on time and context form a higher level of regulatory control known as rheostasis. Rheostasis enables the body to predict and prepare for expected changes in the environment, such as variations in environmental light and temperature, food availability, or behavioural opportunities^11,12^ (**Figure 1c**). A prominent example of such temporal prediction is circadian rhythmicity, which has evolved to align physiological functions with predictable daily cycles. It is now well established that circadian phase has a major impact on diverse physiological processes, including our neurological and cognitive functions^13,14^. This is most clearly exemplified by the sleep-wake cycle^15,16^, in which neural dynamics fundamentally differ between wakefulness and the various sleep stages. To be good regulators, closed-loop DBS systems should therefore be designed with sleep and circadian rhythms in mind^17^ (**Figure 1d**). Feedforward control can be used to change the functional characteristics of sensors, classifiers, control policies and stimulation parameters in the same spirit as rheostasis in natural physiological control.

Parkinson’s disease (PD) is currently the most common indication for DBS therapy^1,2,18^. Sleep disruptions are a common comorbidity exacerbating the disease burden^19^, and ‘standard-of-care’ continuous DBS has been shown to provide benefits for sleep^20–23^. More advanced closed-loop, or adaptive DBS (aDBS) aims to further improve outcomes and reduce side effects by adjusting stimulation in real time, typically using pathological beta-band activity in the basal ganglia as a control signal^24–28^. This approach does not yet factor in large shifts in the dynamic range of beta power linked to the sleep-wake cycle^29–33^ and across sleep stages^34^. While its average power is reduced, persistent nocturnal beta activity is causally linked to Parkinsonian sleep disruption^34–37^. This suggests that aDBS algorithms should account for nocturnal beta dynamics (e.g. adjusting thresholds or setpoints) to avoid potential undertreatment and continued sleep dysfunction at night. Current DBS systems for PD are unable to support such an approach - they have only one mode of operation and are calibrated on daytime brain activity.

Drug-resistant epilepsy is another promising application for DBS. Continuous (or duty-cycled) stimulation of the anterior or centromedian thalamic nucleus (ANT / CMT) is an effective therapy for a proportion of patients^38–41^. However, DBS can disrupt sleep in an amplitude- and frequency-dependent manner^42–44^, while seizures can have a strong association with different sleep stages^45^ and with circadian and other physiological rhythms^46–48^. Interactions between physiological rhythms, seizure risk and DBS therapy settings therefore need to be carefully considered. *Responsive* neuromodulation - delivering stimulation only when abnormal activity is detected - would theoretically maximise sleep quality while minimising seizure duration^49–51^. However, the design of reliable closed-loop control policies is challenging and seizure classifiers will likely perform differently against sleeping vs. waking baseline activity. It is also becoming clear that the seizure-suppressing effect of responsive DBS is at least partially mediated by non-immediate, longer-term effects on neural dynamics^52^ suggesting that some form of baseline stimulation may be beneficial. Notably, different stimulation frequencies may be required for baseline vs. responsive stimulation^53,54^. Therefore, an ideal DBS therapy research platform for epilepsy should be able to leverage feedforward information about patient-specific rhythms to adjust therapy mode, and be able to provide baseline stimulation while monitoring for and responding to breakthrough seizures with different stimulation parameters.

As sleep and circadian disruption is a fundamental underlying component of many neurological and psychiatric conditions^55–57^, improving sleep and circadian rhythms could become a new, explicit aim for DBS therapy. Supporting endogenous rhythmicity is especially critical in applications aimed at restoring arousal and sleep-wake regulation, such as in disorders of consciousness and cognition following traumatic brain injury^58,59^ or in applications targeting autonomic nervous system dysfunction^60,61^. Synchronisation of device actions and physiology can help re-establish a disrupted or absent internal timekeeping signal or worsen it in cases of misalignment^59^.

While it is thus becoming clear that DBS and other bioelectronic systems should account for circadian rhythms and sleep, establishing the optimal course of action for different physiological states requires a clinical-grade platform for therapy research. The field of bioelectronic medicine has recently explored state-responsive therapy^62–64^ proof-of-concept systems, for example through a wireless connection to off-implant data processors^65^. However, this approach is not easily scalable as it places significant burden on patients to manage a complex ecosystem of implantable, wearable and peripheral components.

Here, we introduce the DyNeuMo-2c (Dy2c), the first fully implantable clinical-grade neuromodulation platform to incorporate principles of rheostatic control of healthy human physiology. The Dy2c can switch between multiple closed-loop algorithms based on a scheduled ‘feedforward’ control policy, and enables ‘reactive’ closed-loop therapy based on a neural or motion-based biomarker. Predictive adjustments can be based on *a priori* estimates of therapy need based on disease condition and symptoms, then personalised to account for individual patient chronotypes. The system integrates our previous advances in motion-based adaptive therapy control, (e.g. posture/movement events; DyNeuMo-1)^66^, adaptive therapy algorithms that respond to biopotential input (e.g. local field potentials or LFP; DyNeuMo-2)^67^ and long-term neural biomarker monitoring^68^.

The Dy2c functionality is already set to be enabled as a firmware upgrade for human participants in ongoing DBS trials for post-stroke chronic pain (NCT06387914) and paediatric Lennox-Gastaut epilepsy (NCT05437393). In the periphery, Dy2c is already in use in human participants in trials of pudendal nerve stimulation for urinary incontinence (NCT06885931). This paper presents our system design approach and key validation work in preparation for these studies. Using a state-of-the-art bench testing environment^69^ we demonstrate the ability to implement a therapy schedule optimised for daytime and nighttime phases of typical Parkinsonian brain activity, adjusting the control policy of an adaptive DBS (aDBS) protocol. Next, we demonstrate the Dy2c’s ability to detect epileptic seizure activity while it is providing active baseline stimulation into the medium it is recording from.

Finally, we demonstrate the system capabilities *in vivo* in a healthy non-human primate (NHP) model of daytime sleepiness and vigilance. Here the Dy2c was configured to track and differentially modulate thalamic neuronal activity during sleep, resulting in decreased vigilance and task performance the next day. During the daytime, the Dy2c targeted neuronal signatures of microsleep onset, providing brief enhancements in arousal and restoration of executive attention and performance. The experiment serves to demonstrate the critical importance of circadian phase-alignment between devices and physiology; mismatches of which have recently been reported in human therapy^59,70^. Overall, we believe these cases illustrate the opportunity for a rheostatic device to improve therapies in a broad spectrum of neurological disorders.

## 2. Results

### 2.1 System design

The Dy2c research platform for rheostatic neuromodulation was developed for human clinical application from the outset. It is implemented in the hardware of the cranially mounted Picostim device (Amber Therapeutics, Harwell, UK). The Dy2c can utilise a biopotential amplifier as well as an accelerometer for sensing neural biomarkers and estimating posture or behavioural state and implementing closed-loop therapy via an on-board processor, and is integrated in a multi-level pipeline for therapy development through external connectivity (**Figure 2a**). At the time of writing, the Picostim hardware has been implanted in human clinical trials for a range of neurological disorders, including Parkinson’s disease (NCT03837314), multiple system atrophy (NCT05197816), Lennox-Gastaut Syndrome epilepsy (NCT05437393), and chronic pain (NCT06387914), as well as in the periphery of the central nervous system in mixed urinary incontinence (NCT06885931). The core innovations of the Dy2c firmware enable rheostatic operation and chronotherapy with existing implants. The Dy2c design adhered to EN 60601-1-10 standards for physiological closed-loop controllers, EN ISO 14971:2019+A11:2021 for risk management, EN 62304:2006+A1:2015 class C for the software and firmware. The overall system was developed under an EN ISO 13485:2016-compliant quality management system for human medical devices.

**Figure 2:**
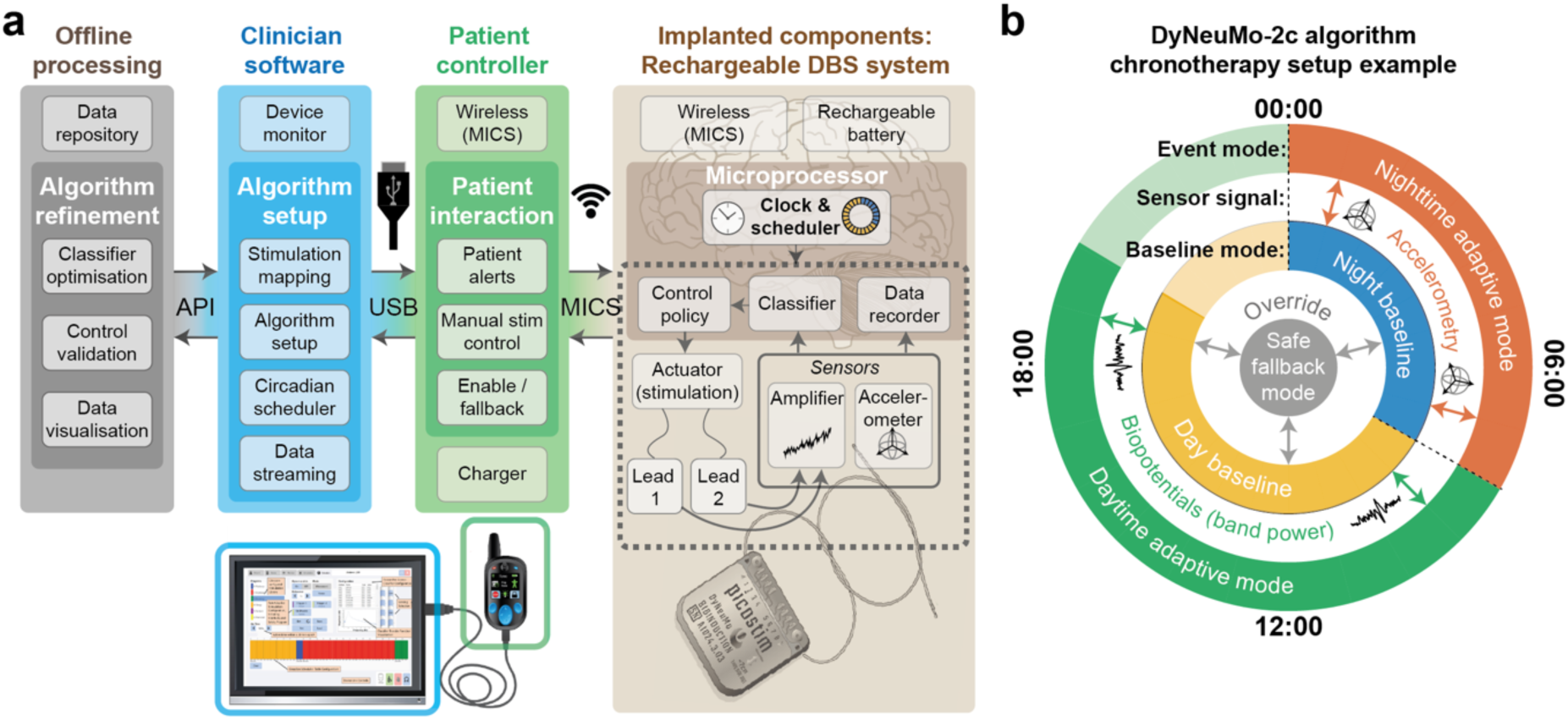
Dy2c as a platform for the research of rheostatic control in medical devices. **a)** Dy2c technology stack showing how the Dy2c operates as a platform for rheostatic therapy development, highlighting implanted elements as well as essential functions of the patient controller, clinician software on tablet, and offline processing. Human brain adapted from original by John Chilton (https://doi.org/10.5281/zenodo.3925988). **b)** An example of how the functionality of the Dy2c can be used to set up a dual-profile chronotherapy algorithm. In this example, there are two different profiles active at different times of day, which can be selected based on the patient’s sleep routine. During the patient’s waking hours (e.g. 07:00–00:00) the ‘daytime baseline’ stimulation is provided, with an adaptive change to stimulation implemented in response to variations in a feature extracted from biopotentials (LFPs) recorded from a particular electrode montage relative to a patient-specific and time-dependent threshold. During the patient’s normal sleeping hours (e.g. 00:00–07:00), the system switches to a different algorithm configuration profile with night- and sleep-optimised baseline stimulation parameters, and implements a different adaptive change to stimulation if a feature is detected on the accelerometer data stream (e.g. triggered when a patient needs to get up during the night).

For rheostatic therapy implementation, a hierarchical scheduler controls switches between different system configuration ‘profiles’ according to time of day (**Figure 2b**). Profiles have independent baseline effectors (stimulation amplitudes, frequencies and electrode contacts), independent sensing montages, independent classifier parameters, and independent control policies for automated therapy changes in response to classifier outputs. To help identify therapeutically relevant settings for the circadian scheduler, the Dy2c supports a resolution looping data recorder (‘loop-recorder’) with configurable time resolution for tracking time-dependent signal characteristics. For example, average LFP band-power may be logged at a rate of 0.5 Hz for over 30 hours to help establish patient-specific circadian patterns (i.e. chronotype). Temporal resolution can be traded for recording length, potentially supporting weeks to months of storage to capture infradian (slower than daily) rhythms. For proof-of-concept research implementations of advanced stimulation patterns or closed-loop approaches, the system can stream data, and update stimulation parameters in real time via an application programming interface. To facilitate research and minimise surgical risk, the Dy2c firmware can be made available through a wireless upgrade to any previously implanted Picostim device, providing a rapid route to follow-up clinical studies without further surgical burden on patients.

### 2.2 Bench verification: Scheduled algorithm profile switches for day- and night-optimised closed-loop therapy in PD

For rapid simulation of clinical use cases, we used the NeuroTest platform: a custom benchtop system for validation of adaptive sensing and stimulation systems^68,69,71^ (**Figure 3a**). This platform enabled us to accurately reproduce a pre-recorded neural signal as an electric field in a saline medium for the Dy2c to record, and to stimulate into the same medium^69^. This allows the stimulation pulse and injected neural signal to interfere (as they would *in vivo*), resulting in realistic evaluation of sensing performance during active stimulation.

**Figure 3:**
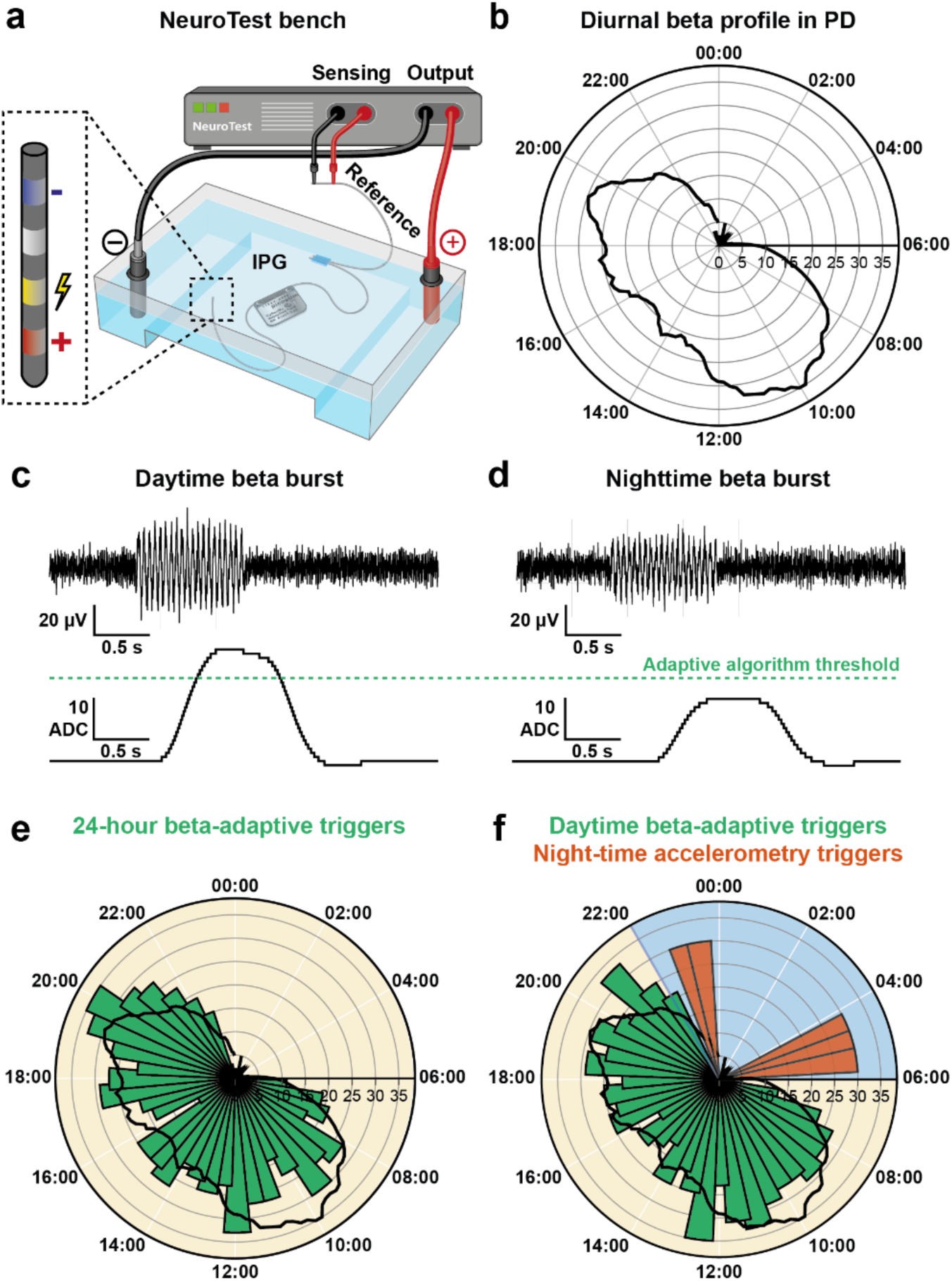
NeuroTest bench demonstration of time-of-day-based switch between beta-adaptive and motion-adaptive therapy for PD. **a)** Test bench setup using the NeuroTest platform. The NeuroTest system injects pre-recorded LFP signals into a saline tank in which the Dy2c is immersed. The Dy2c registers this activity using the simultaneous recording and stimulation montage illustrated in the inset. **b)** Daily profile of subthalamic nucleus beta-band microvolt power for a typical PD patient^29^. **c)** Simulated subthalamic nucleus LFP with beta burst at daytime level of beta power (top), with beta power extracted by the closed-loop algorithm and the daytime algorithm threshold (bottom). Note that the beta envelope crosses the adaptive algorithm threshold at daytime level. **d)** Simulated LFP with beta burst at nighttime level of beta power (top), with beta power feature extracted by the closed-loop algorithm and the adaptive algorithm threshold (bottom). Note that this nighttime level beta burst does not cross the adaptive algorithm threshold. **e)** Histogram of adaptive algorithm triggers when streaming 24-h of simulated data with beta bursts modulated according to time of day into the virtual brain test setup with Dy2c running adaptive therapy based on a daytime-optimised threshold. Note the adaptive algorithm shows few to no triggers between 22:00 and 06:00. **f)** Histogram of adaptive algorithm triggers in response to the same test signal, with the Dy2c scheduled to change between a daytime (06:00–22:00) and nighttime profile. The daytime profile used the aDBS algorithm with an optimised threshold, while the nighttime profile was based on constant baseline therapy with stimulation increases triggered by motion events. Note a similar number of daytime adaptive therapy and nighttime motion-triggers when a researcher generated motion triggers by physically moving the device during the night.

In PD, the only currently approved closed-loop therapy (aDBS^28^) seeks to suppress subthalamic nucleus (STN) LFP activity in the beta frequency range (∼13-30 Hz) which is associated with symptom severity. To investigate the predicted behaviour of aDBS for PD across the 24-hour cycle, simulated Parkinsonian STN LFPs were played into the NeuroTest setup. LFPs contained pseudorandom patterns of simulated beta bursts (peak frequency 20 Hz) that were modulated according to the diurnal profile reported from chronic 24-h recordings in a PD patient (**Figure 3b-d**)^29^. First, we investigated the behaviour of a single-threshold beta-adaptive algorithm with a threshold determined according to the daytime beta power range, with a baseline bilateral stimulation at 0.5 mA (126 Hz, 90 μs pulse width), increasing to 3 mA when the beta signal exceeded the threshold. This resulted in the expected adaptive algorithm triggers during daytime, but no adaptive triggers during the night when beta activity is low due to sleep (**Figure 3e**).

Next, using the Dy2c’s scheduler, we set up a time-of-day-dependent dual-profile adaptive programme. In the simplest embodiment, we used a beta adaptive algorithm during the day (6:00–22:00) and a nighttime baseline stimulation (i.e. open-loop stimulation) with an accelerometer-based motion adaptive mode (22:00–6:00) to address spontaneous awakenings. The nighttime profile was designed to have a slightly higher baseline amplitude (1 mA) hypothesised to be more protective of sleep, with a stimulation increase (to 3 mA) for accelerometer-based events representing patient activity such as getting out of bed to use the toilet (simulated by a researcher physically moving the device within the test setup during two nighttime episodes). This resulted in a similar profile of sense adaptive algorithm triggers during the daytime (6:00–22:00), but scheduled transition to a nighttime stimulation baseline and the expected motion-adaptive triggers during the night (22:00–06:00; **Figure 3f**).

### 2.3 Bench verification: Adaptive therapy for epilepsy

In DBS for epilepsy, current practice is either to provide stimulation the same (continuous or duty-cycled) frequency and amplitude^40,41^, or responsive stimulation where stimulation is only delivered in response to detection of abnormal brain activity^49–51^. No DBS devices currently align open- or closed-loop therapy modes with patient-specific diurnal seizure rhythmicity. Here we demonstrate the ability to deliver *adaptive* DBS for epilepsy – implementing CMT LFP-based seizure detection during active stimulation, and a switch to different stimulation settings upon such detection.

LFP seizure events from the CMT of a patient with genetic generalised epilepsy were streamed into the NeuroTest setup and recorded by the Dy2c (**Figure 4a**). The envelope of the frequency band between 10–20 Hz was found to be a feature suitable for the classification of seizure occurrence and resulted in classifier with a receiver operating characteristic (ROC) area under curve (AUC) of > 0.95 (0.995) and a F1 score of 0.87 over a 24-hour cycle^68^. To demonstrate the robustness of the classifier to interference from active stimulation, the same data were re-played while the Dy2c was running a closed-loop algorithm with a baseline low frequency (6 Hz) bilateral stimulation (1 mA, 6 Hz, 100 μs pulse width), and adaptive switches to high frequency bilateral stimulation (4 mA, 126 Hz, 100 μs) upon seizure detection (**Figure 4b**).

**Figure 4:**
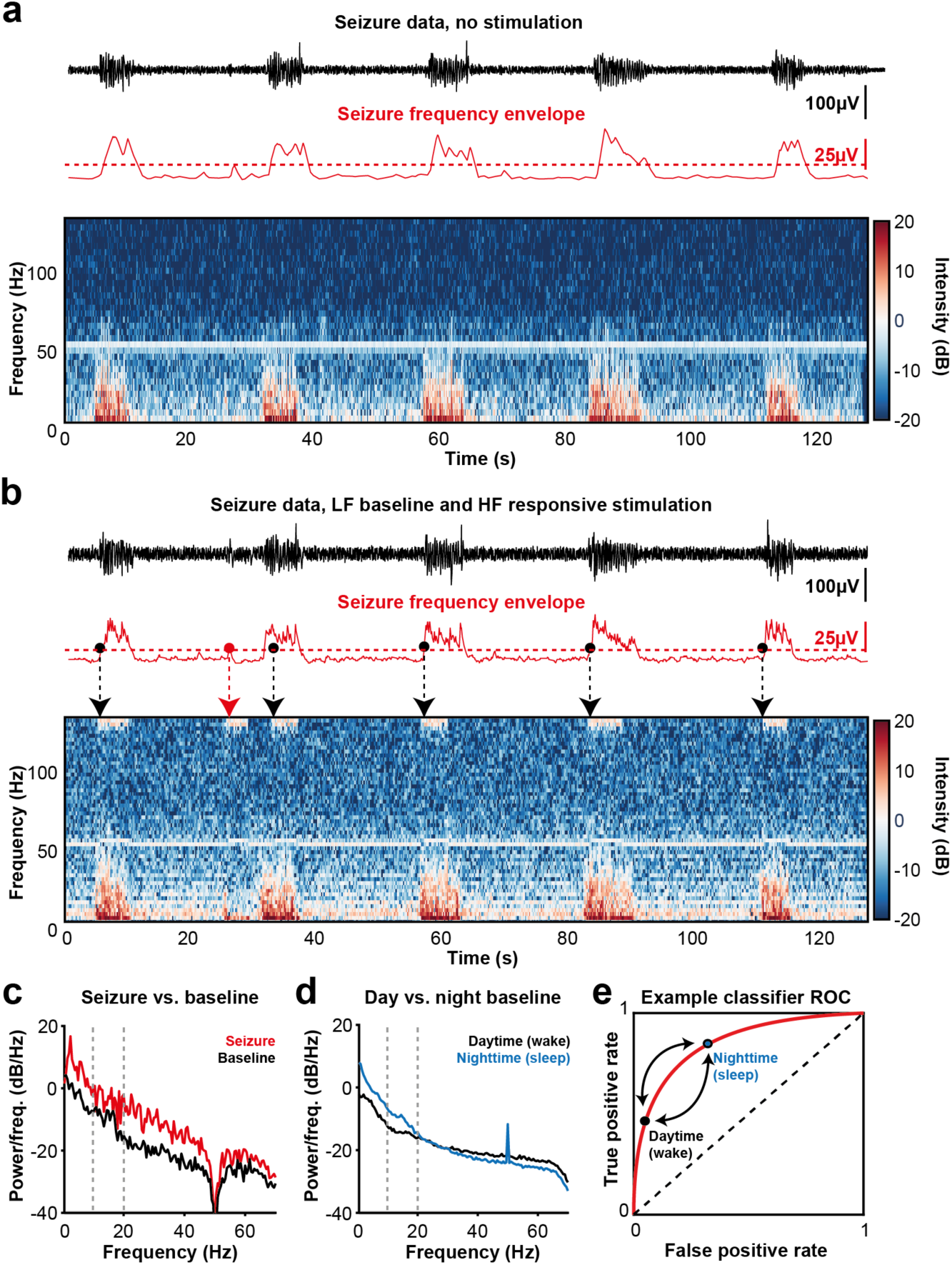
The Dy2c supports seizure classification in the presence of different stimulation frequencies, and allows optimised therapy settings for day- and nighttime use in epilepsy. **a)** LFP of seizure events from a generalised epilepsy patient concatenated together at 10 s intervals and streamed into the NeuroTest and recorded by the Dy2c (top, black), envelope of seizure frequencies of interest (10–20 Hz, middle, red), and spectrogram of this LFP data as recorded by the Dy2c (bottom). **b)** Same seizure data, envelope and spectrogram as in a), but recorded by the Dy2c during baseline stimulation at 6 Hz with adaptive stimulation triggers at 126 Hz. Note the stimulation artifact bands appearing in the spectrogram at 126 Hz (bottom) upon seizure detection. **c)** Power spectrum of CMT LFP activity at (daytime) baseline (10 s of baseline activity in the pre-seizure period) and during a (daytime) seizure (the 10 s following seizure onset). During the seizure event there is a broadband increase in power compared to baseline, non-seizure activity. Dotted lines represent the seizure-diagnostic frequency band across the data set (10-20 Hz). **d)** Power spectrum of non-seizure CMT LFP activity during 10 minutes of daytime wakefulness and 10 minutes of nighttime sleep, illustrating pronounced shifts in baseline power across frequency bands that overlap with the putative seizure-diagnostic band (10-20 Hz). **e)** Example receiver operating characteristic (ROC) curve for Dy2c seizure detector for daytime and nighttime. Points on the ROC curve indicate proposed bias for day and nighttime adaptive algorithm use. During daytime operation, it may be desirable to bias the classifier to minimise false positives to prioritise the delivery of open-loop therapy during daytime operation. In contrast, if poor therapeutic performance is reported for open-loop therapy at night-time the classifier can be biased to greater sensitivity (i.e. increase the false positive rate of the classifier) so that adaptive delivery of DBS therapy can be delivered more at nighttime.

The Dy2c maintained sensing performance and the envelope of the 10-20 Hz band remained highly predictive of seizure occurrence. Classifier performance was only modestly affected by active stimulation (ROC AUC reduced from 0.995 to 0.922), a performance decrement unlikely to affect clinical utility.

Additionally, classifier parameters may benefit from differential operation during the day and night. **Figure 4c** shows the spectral characteristics of the CMT LFP during an example seizure compared to baseline, highlighting an increase in power across a broad range of frequencies. While the 10-20Hz band was most diagnostic for seizure detection across this data set, **Figure 4d** illustrates pronounced shifts in baseline CMT LFP activity in this frequency band between daytime and nighttime sleep. Therefore, the optimal classifier parameters for adaptive DBS for epilepsy may vary according to time of day. Such diurnal adjustment of the classifier performance is possible with the Dy2c scheduler and is illustrated in **Figure 4e** where daytime and nighttime operation of the classifier could correspond to biasing the algorithm to different points on the receiver-operator characteristic (ROC) curve.

### 2.4 *In vivo* validation in a model of dysfunctional executive attention

#### 2.4.1 System implantation

To characterise the *in vivo* performance of Dy2c and highlight its potential as a platform for advanced preclinical rheostatic therapy research we employed it in a NHP vigilance modulation paradigm. A recent human feasibility study aimed at treating enduring cognitive dysfunction in moderate-to-severe traumatic brain injury patients^59^ revealed an unmet need to address sleep and circadian disturbances. As a first step towards developing this system, we implanted a healthy NHP (**Figure 5a, Supplementary Figure 1**) to identify neurophysiological biomarkers linked to fluctuations in arousal and executive attention throughout the day and night. We then used these biomarkers to validate the streaming, recording, and aDBS capabilities of the Dy2c using scheduled daytime and nighttime aDBS.

**Figure 5:**
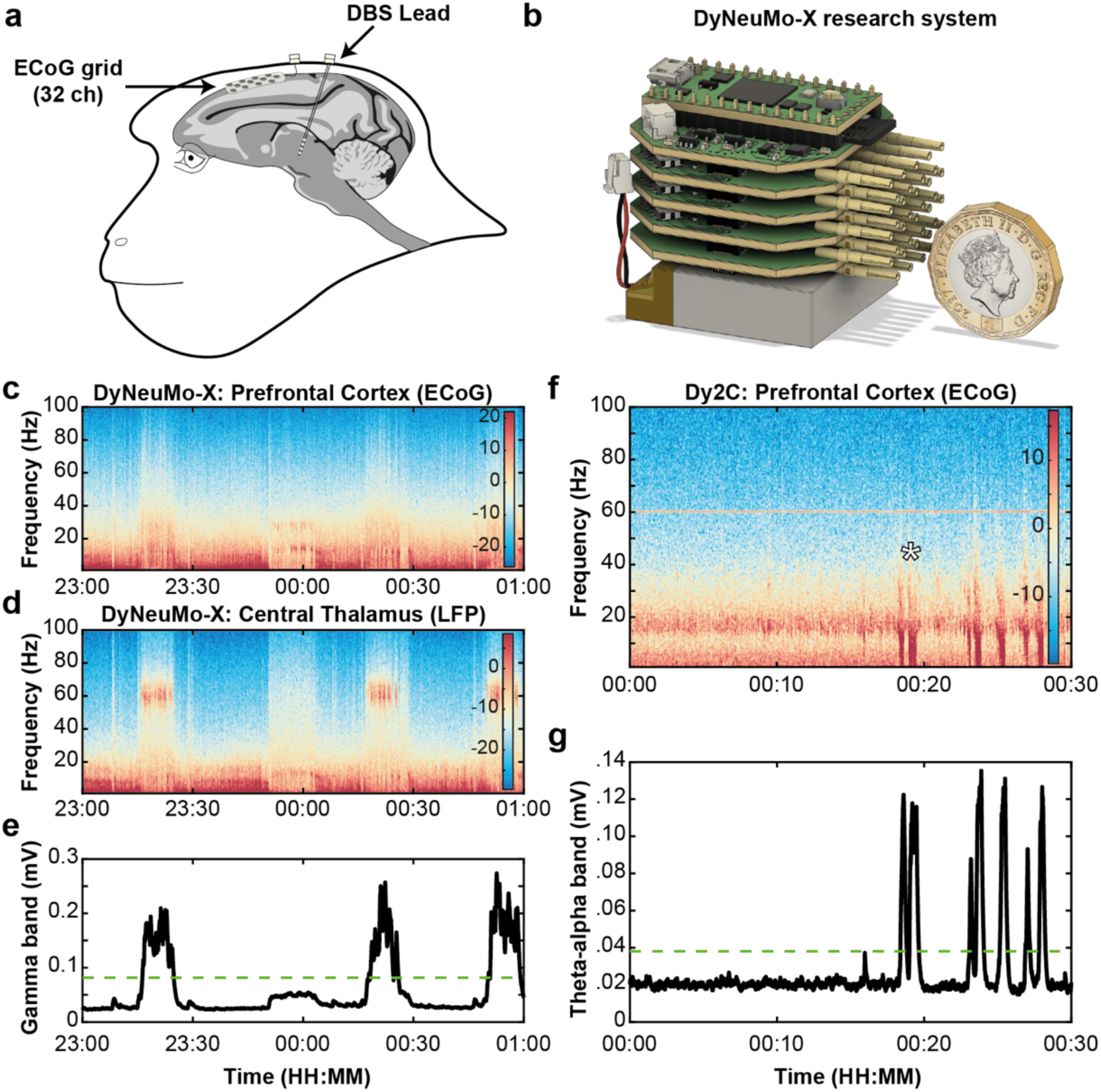
NHP vigilance task performance and nighttime aDBS. **a)** An illustration of the animal’s cephalic implant that included a multichannel ECoG array placed over the prefrontal and premotor cortices and DBS leads placed within the central thalamus, for each hemisphere. **b)** The DyNeuMo-X research system consists of a base data collection board, up to three sensing and one stimulation boards (8-channels each), connected to a Lithium-Polymer battery for long duration (> 20 hours) experiments. A 3D printed case secured the system within the implant’s protective cap (**Supplementary Figure 1d**). **c)** Spectrogram of prefrontal ECoG activity recorded over right cortical areas 8AD and 6DC during the set 18:00 – 06:00 lights off period. The animal woke up and moved about its home environment at 23:51 and then resumed sleep. Estimated with 5 s windows, 20% overlap, with 0.5 Hz resolution. The colour scale is expressed as the log power on a microvolt signal. **d)** Spectrogram of LFP activity recorded within the right central thalamus, simultaneously with c). This segment illustrates the repeating periods of enhanced gamma-band activity. **e)** Output of a software emulation of the Dy2c system’s signal processing chain, tracking the envelope magnitude of the gamma-band activity (55-65 Hz) in the thalamic LFP signal shown in D. The dotted green line at 80 μV shows the threshold level used to trigger aDBS in subsequent experiments. **f)** A spectrogram of prefrontal cortical ECoG activity (same areas as in c)) streamed with the Dy2c system while the animal performed the vigilance task. The white asterisk highlights two consecutive periods of drowsiness and full eye closure. **g)** A software emulation of the Dy2c system’s signal processing chain, tracking the envelope magnitude of the theta-alpha band (5–10 Hz) in the cortical ECoG signal shown in F. The dotted green line at 38 μV shows the threshold level used to trigger aDBS in subsequent experiments.

We first used a custom research device (**Figure 5b, Supplementary Figure 1d**), the DyNeuMo-X, to identify the neurophysiological biomarkers linked to fluctuations in arousal and executive attention. Compared to the Dy2c, the DyNeuMo-X supports higher channel counts with higher temporal resolution. Accordingly, the DyNeuMo-X was used to concurrently record prefrontal ECoG (**Figure 5c**) and central thalamic LFP (**Figure 5d**) during naturalistic conditions and sleep. During sleep, we observed spectral features consistent with established sleep stages, such as repeated bouts of slow wave activity that included sleep spindles (9–16 Hz) and pronounced low frequency activity (0.5–4 Hz), characteristic of N2 and N3 sleep. We also identified a unique and robust spectral feature within the thalamic LFP that exhibited repeating epochs of enhanced ‘gamma-band’ (55–65 Hz) activity (**Figure 5d, Supplementary Figure 2b,e**). This feature was present bilaterally but was most prominent within the right rostral DBS lead (**Supplementary Figure 1c, 2b,e**). Video and motion capture confirmed that the animal was sleeping during these gamma-band epochs (n=14), lasting on average 7.5 min (1.3–12.3 min). A classifier based on the gamma-band activity (55–65 Hz) (**Figure 5e, Supplementary Figure 2c**) was uploaded to the Dy2c to track nighttime gamma activity (**Figure 6c–f**). These recordings (**Supplementary Figure 2f**) confirmed the regularity of the gamma-band epochs, here lasting on average 8.1 min (2.9– 11.9 min), and established the 80 μV threshold to trigger aDBS in subsequent experiments.

**Figure 6:**
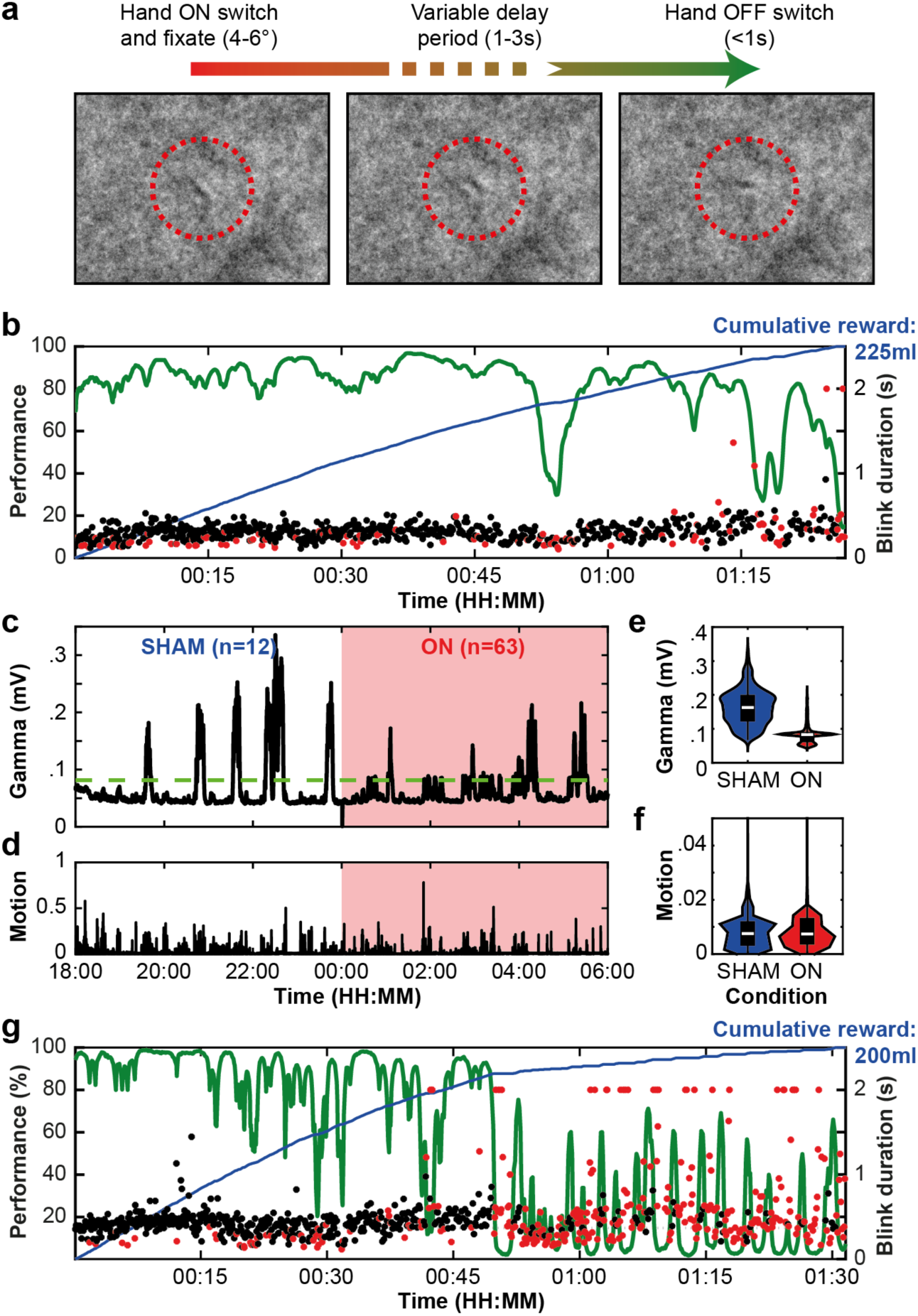
Closed-loop targeting of nighttime gamma bursts results in reduced vigilance the next day. **a)** Example of the vigilance task. The visual field consisted of a centrally placed Gábor-patch over noise. In self-initiated trials, the patch’s orientation would randomly change, indicating a chance to obtain a liquid reward. **b)** The animal’s unperturbed performance on the task. The green curve represents a smooth estimate of performance. Pupillometry was used to identify the time and duration of eye blinks. Blinks that occurred during correct trials are indicated by black dots and during incorrect trials by red dots. The blue trace represents the accumulating liquid reward across the trial (total reward for the day’s session was 225 ml). **c)** Overnight recording of gamma-band activity (55–65 Hz) from the right DBS lead using the Dy2c. The dotted green line at 80 μV represents the aDBS activation threshold, scheduled to be active (ON) in the second half of the night (00:00–06:00). When the envelope magnitude of the gamma-band signal exceeded the threshold and aDBS was active, stimulation was triggered in the right DBS lead (ramped from 0.1–2.5 mA). **d)** Overall motion during the nighttime hours was derived from the 3-axis accelerometer synchronised to the Dy2c. Values were normalised to the peak motion recorded, representing daytime activity. **e)** Violin plots of the gamma-band (55–65 Hz) signal shown in b), restricted to when aDBS was triggered during the ON (00:00–06:00) and SHAM (18:00–00:00) stimulation periods. The width of the violin plots represents the probability density of the magnitude values on the vertical axis. The black box plot shows 25th and 75th percentile for each distribution, and the white line is the median value. **f)** Violin plots of the motion values during the same periods in e). **g)** The animal’s performance of the vigilance task the day following the 00:00–06:00 aDBS experiment shown in c). Annotations are the same as in b).

Validation of the Dy2c system was then conducted with the NHP under three conditions: 1) free behaviour and sleep within a restricted region of the animal’s home environment for video monitoring, 2) behavioural performance during a visuomotor reaction time task (i.e. vigilance task) lasting 1–2 hours with liquid rewards, in which the animal was head-restrained, and 3) free behaviour and sleep within the home environment that included interactions with its roommate.

#### 2.4.2 Disruption of nighttime gamma-band epochs disrupts performance the next day

In a clinical feasibility study^59^ daytime therapy was inadvertently applied in one patient throughout the night, which markedly impacted sleep and exacerbated daytime symptoms. To model this clinical finding, we used nighttime aDBS and to explore its impact on next day performance (Figure 6). In a previous study^72^, we used daytime aDBS to restore an animal’s performance on the vigilance task (**Figure 6a**) and here we extended this work using nighttime aDBS. When motivated, animals typically perform the vigilance task well (80–100%) at the start of a session (during the first 30–45 min) and then gradually reduce their engagement as satiety, boredom and drowsiness increase. We hypothesised that the animal’s performance would be negatively impacted by disrupting the night-time bursts of gamma activity with closed-loop DBS. In the day prior to the nighttime aDBS (**Figure 6b**), the animal’s performance was good (80– 100% correct) until about 50 min into the session, when performance first dropped below 50%. At 01:15 the animal’s blink durations started to increase, and performance began to vary. At the end of the session full eye closures (lasting >1 s) marked the onset of drowsiness, at which time the animal ceased to engage with the task. These results are largely consistent with our prior studies in animals performing similar tasks^73–75^. The Dy2c device was then secured to the animal and scheduled to trigger aDBS during the night only (00:00–06:00) using the gamma-band filter (55–65 Hz) and 80 uV threshold.

In **Figure 6c-f** we describe the results of the scheduled nighttime aDBS. In the first half (18:00– 00:00), the Dy2c was programmed to deliver SHAM stimulation (at subthreshold level 0.1 mA), while logging all threshold crossings. In the second half of the night (00:00–06:00) stimulation was triggered when the gamma-band signal crossed the threshold. In the first four hours (00:00– 04:00), stimulation consistently reduced thalamic gamma power, while during the last two hours (04:00–06:00), the gamma-band activity rebounded to levels observed in the SHAM period, despite ongoing aDBS. Overall, the aDBS ON protocol, when applied during the second half of the night, did significantly reduce the thalamic gamma-band signal (p < 0.0001, Mann-Whitney U-test) despite the rebound periods, when compared to SHAM stimulation during the first half of the night (**Figure 6e**). Of note, the animal’s motion was unaffected during aDBS (p > 0.9999, Mann-Whitney U-test).

As hypothesised, the animal’s performance diminished the next day, consistent with reduced vigilance following aDBS-induced sleep disruption during the previous night (**Figure 6g**). Long blinks and eye closures occurred early in the session (within the first 15 min), and there were extended periods of drowsiness and putative ‘microsleep’ periods (eye closures longer than 1 s) in the second half of the session. This result was reproduced the following week (**Supplementary Figure 4**).

#### 2.4.3 Disruption of daytime low frequency activity restores daytime vigilance

We next implemented an aDBS protocol designed to promote vigilance during daytime hours (06:00–18:00) while the animal performed the vigilance task (**Figure 7a,b**) and while it freely behaved in its home environment (**Figure 7c,d**). We targeted cortical theta-alpha band (5-10 Hz) activity, linked to fluctuations in arousal and executive attention that impact performance^72^. First, a computer emulation of the low frequency theta-alpha signal (5–10 Hz), recorded from the prefrontal cortex by the DyNeuMo-X (**Figure 5f,g**), was used to develop a classifier to trigger aDBS when the animal closed its eyes for extended periods of time (**Supplementary Figure 3**). In this session, the Dy2c was programmed to deliver aDBS (10 s debounce, ramping from 0.1 mA to 4.0 mA), on detection of cortical theta-alpha (**Figure 7a,b**). As expected, long blink durations and full eye closures preceded each aDBS period, lasting on average 0.75 s (ranging 0.19–7.86 s). Blink durations following aDBS were markedly shorter (mean 0.24 s, ranging 0.10–1.1 s), reflecting an overall increase in arousal. Of note, the animal resumed performance of the task after 18 of the 32 aDBS periods. These preliminary results suggest that aDBS could be used to transiently enhance arousal to restore executive attention during increasing mental fatigue.

**Figure 7:**
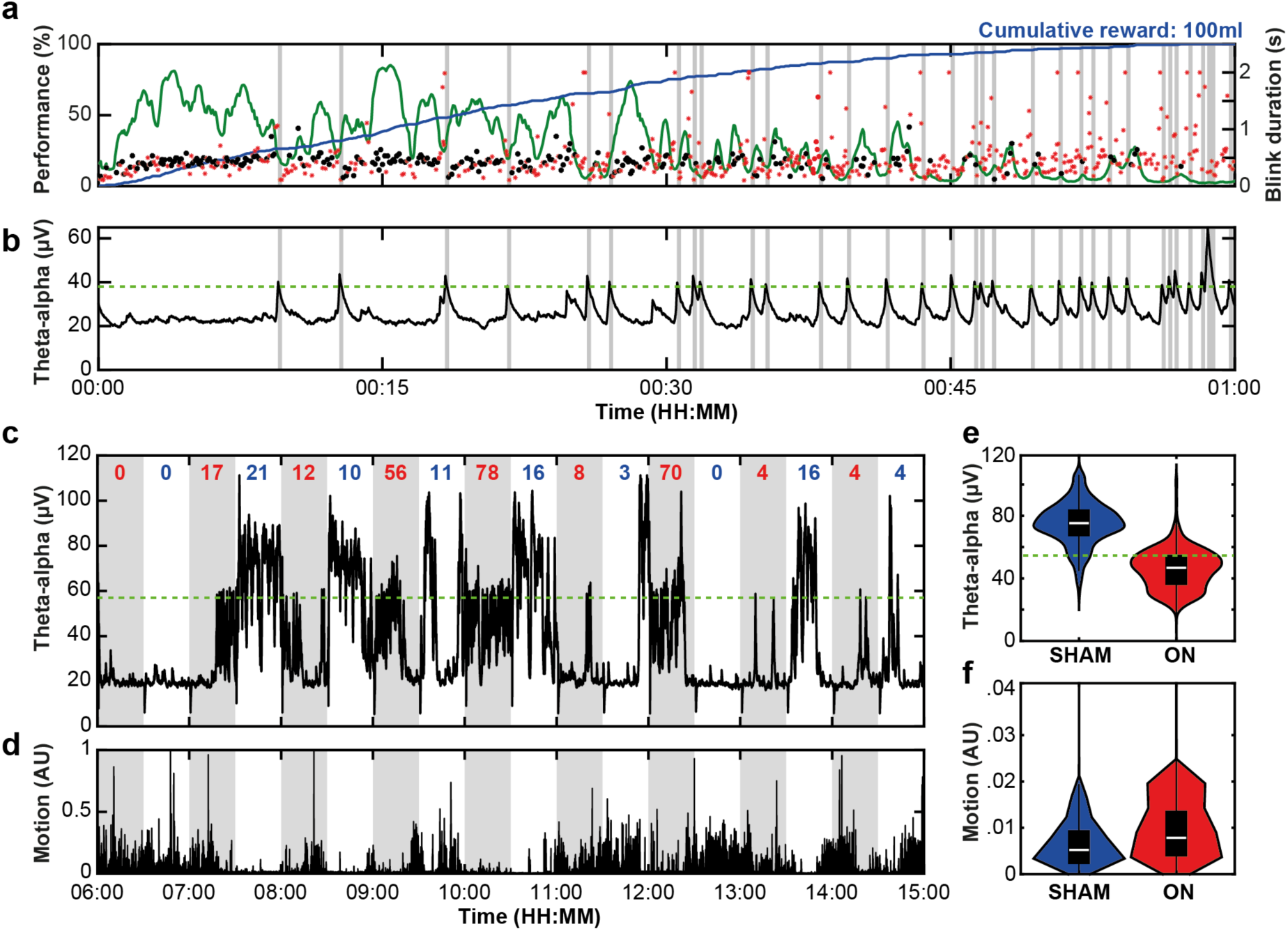
Theta-alpha activity (5-10 Hz) signals a decline in performance, stimulation can extend vigilance. **a)** The animal’s performance on the vigilance task, but while it was not restricted from access to water in the home environment and instead rewarded with 100% apple juice to provide motivation to engage the task. Blinks that occurred during correct trials are indicated by black dots and during incorrect trials by red dots. Blinks and eye closures longer than 2 s are set to a ceiling value of 2 s. The blue trace represents the accumulating juice reward across the session (total juice rewarded was 100 ml). Periods of aDBS activation shown as vertical grey bars. **b)** While the animal performed the vigilance task for apple juice rewards, an aDBS protocol was used by the Dy2c to continuously record theta-alpha activity (5–10 Hz) from the right ECoG contacts placed over areas 8AD and 6CD. The dotted green line at 38 μV represents the envelope magnitude of the signal that triggered the aDBS algorithm. Periods of stimulation (ramped from 0.1–4.0 mA) are shown as vertical grey bars. **c)** While the animal moved freely throughout his home environment the Dy2c continuously recorded theta-alpha band activity (5–10 Hz) from the right ECoG contacts placed over areas 8AD and 6CD. The dotted green line at 57 μV represents the envelope magnitude threshold on the in-band activity, that would trigger a change in stimulation. Stimulation increased (ramped from 0.1–4.0 mA) during the ON phase (06:00–14:30) shown in grey. Intervals in white represented SHAM phases, with triggers activating a sub-threshold programme. The number of threshold crossings in the ON and SHAM phases are shown in red and blue, respectively. **d)** Overall motion during the nighttime hours was derived from the 3-axis accelerometer synchronised to the Dy2c. Values were normalised to the peak motion recorded, representing daytime activity. **e)** Violin plots of the theta-alpha signal (5–10 Hz) shown in c) at points of aDBS was triggering during the ON (red) and SHAM (blue) periods. The width of the violin plots represents the probability density for each voltage value. The black box plot shows 25th and 75th percentile for each distribution, and the white line is the median value. **f)** Violin plots of the motion values during the same periods in e).

The final validation experiment was conducted using the same daytime theta-alpha classifier to trigger aDBS while the animal freely moved about its home environment and interacted with its roommate. Of note, the aDBS threshold was increased (from 38 μV to 57 μV) to account for elevated theta-alpha activity associated with extended eye closures lasting > 1 s. Here the 24-hour circadian scheduler was used to toggle between the aDBS ON protocol and the investigator programmed SHAM protocol, every 30 min (**Figure 7c,d**). The device logged the number of threshold crossings during each 30 min period. In this 9-hour session, a total of 249 threshold crossings occurred during the aDBS ON periods and 81 crossings during the SHAM periods. When comparing the distribution of theta-alpha intensity (**Figure 7e**) and overall motion (**Figure 7f**) across conditions, aDBS markedly reduced the cortical theta-alpha band activity (p < 0.0001, Mann-Whitney U-test) and modestly but significantly increased the animal’s overall movement (p < 0.0001, Mann-Whitney U-test). An additional 12-hour session (**Supplementary Figure 5**) replicate this result, thus demonstrating the Dy2c’s capacity to reliably modulate intrinsic fluctuations in daytime arousal under more naturalistic conditions.

## 3. Discussion

### 3.1 Achieving rheostatic control with bioelectronic systems

The Dy2c is a highly configurable, fully implantable, clinical grade platform for research, development and implementation of rheostatic closed-loop neuromodulation therapy. We highlighted the system’s ability to implement scheduled changes between closed-loop algorithms, and to maintain biopotential sensing and classification performance during different active baseline stimulation frequencies using bench tests relevant to clinical treatment of PD and epilepsy. Next, in a series of *in vivo* experiments in a healthy NHP, we demonstrated safe and effective modulation of an ultradian fluctuation in arousal during sleep, and modulation of arousal and behavioural performance during the day. The ability to implement the Dy2c firmware set on previously implanted Picostim systems, as well as streaming support for off-device processing, provide a rapid potential route to advanced therapy research in human participants.

### 3.2 Benchtop demonstration of novel device capabilities

The Dy2c enables pre-emptive adjustments to stimulation, allowing anticipation of patients’ specific symptom or biomarker chronotypes. Our benchtop implementation of scheduled aDBS for PD highlights the power of time-contingent profile switches in the context of daily fluctuations in the beta control signal^29–33^, which are not captured by current aDBS algorithms^24–28^. The freedom to switch between daytime and nighttime algorithms allows these to be optimised to support wakefulness and sleep, respectively. In this case, the sleep-optimised algorithm utilised accelerometer input to respond to night-time awakenings^66^. Another approach would be to refine the nighttime classifier with a lower threshold to make the detector more sensitive to nocturnal beta bursts, or adjust the nighttime classifier towards higher beta frequencies that are most correlated with sleep disruption^34^ and have less overlap with the frequency range of sleep spindles^76–78^.

Our epilepsy-focused bench test using streamed intracranial seizure data demonstrates the Dy2c’s ability to provide closed-loop stimulation to control breakthrough seizures during active baseline stimulation, an advantage over current responsive therapy systems^49–51^. The ability to operate a closed-loop algorithm while providing *low-frequency* stimulation also improves significantly on the two currently FDA-approved adaptive therapy systems. The Medtronic Percept only supports adaptive sensing at stimulation frequencies above 40 Hz^79^, while the Neuropace RNS does not allow for concurrent sensing and stimulation at all^49–51^. Neither device supports rheostatic configurability of setpoints for tuning algorithm parameters in synchrony with a patient’s symptom-chronotype or sleep-wake cycle. While a commercial vagal nerve stimulator for epilepsy supports scheduled diurnal adjustments^80^, this is to reduce the side effect of sleep apnea, not a component of primary therapy.

These results are not exhaustive of the capabilities of the Dy2c but are intended to showcase a feature-set with near-term relevance in clinical practice.

### 3.3 *In vivo* validation in healthy NHP

Our experimental manipulation of a healthy NHPs performance of a vigilance task highlights the ability of the Dy2c to effectively operate on multiple timescales *in vivo*. This functionality allowed the detection unique different neural features (e.g. high gamma-band or theta-alpha activity) linked to specific states of arousal at different times of day, to modulate local and global neuronal activity based on the occurrence of these features and to utilise this stimulation to reliably achieve bidirectional impacts on sleep/vigilance and behaviour. Moreover, this approach could be used to exploit rheostatic principles by scheduling the reactive closed-loop modulation to sleep or wakefulness. These results provide a foundation to develop new bidirectional systems that can address sleep and circadian disturbances in TBI and other conditions.

The observed effects of DBS on vigilance reinforced the need for synchronisation of device operation with intrinsic physiology when deployed in patients. Nocturnal aDBS, upon detection of sleep-associated gamma-band activity, resulted in robust suppression of this neuronal feature, adversely impacting next day performance. On the contrary, properly timed daytime suppression of the 5–10 Hz theta-alpha activity in the prefrontal cortex promoted real-time improvements in vigilance and restoration of performance. The ability of the Dy2c to precisely perturb sleep phases and other physiological rhythms makes it a promising research tool for circadian and sleep neuroscience. For example, a series of non-pharmacological sleep disruptions in healthy NHPs could be used to induce excessive daytime sleepiness (EDS) and dysregulation of arousal and vigilance, as a model of the cognitive dysfunction in traumatic brain injury patients^58,59^.

The discrepancy between the animal’s level of arousal and our ability to capture its fluctuations via motion detection is a limitation of this study. The animal was not instrumented with EOG, EMG, ECG, nor occipital EEG/ECoG channels to permit full sleep stage classification per gold-standard polysomnography guidelines^81^. Future studies focused on the neuroscience of sleep would need to apply more advanced instrumentation for full validation.

### 3.4 Device limitations

The Dy2c is a proof-of-concept for rheostasis-like behaviour in an active medical device through hierarchical operation of a (predictive) scheduler and (reactive) closed-loop functionality. The intent is to motivate device designers to consider hierarchical control with rheostasis in their future designs. The current system has limitations given the constraints of implementation on human clinical-grade implantable hardware (the Picostim). For example, Dy2c is limited to monitoring a single biopotential signal at any one time – though this can be multiplexed across all electrode combinations and the montage includes unique cross-lead sensing capability even in the presence of stimulation. However, stimulation electrodes cannot be used for sensing within the same scheduler epoch. The current embodiment only allows two scheduled configuration profiles, but this was chosen consciously to limit the degrees of freedom in device programming, which is already complex in adaptive systems. The biopotential classifier implements a single-threshold aDBS algorithm, chosen to replicate the seminal design explored in PD^27^; dual threshold designs or other approaches are feasible but were not explored in detail. The Dy2c has to manage finite onboard memory, which motivated the design of a *rolling loop* recorder to gather data for classifier training.

The ability to optimise neuromodulation therapy to safeguard or improve sleep is one of the most important motivators for implementing different therapy profiles, but on the Dy2c this is currently approximated through changing configuration according to patient-specific symptom-chronotypes. Dy2c does not currently gate algorithmic operation based on detected brain state, unlike recent work using a wirelessly connected system with offline processing^63,64^. This means that ‘brain-state’ dependent therapy currently is implemented via a feedforward expectation of the patient’s sleep-wake rhythm (chronotype), which does not account for variability across days.

However, it should be noted that the *prescriptive* nature of the scheduler may be an advantage in some use cases – especially to reinforce desired patterns of physiology or proactively avoid symptoms during critical time windows. For example, when stimulation is adjusted for nighttime to prevent sleep disruption, the ideal system behaviour is to change stimulation parameters when the patient *intends* to fall asleep rather than after sleep is already detected, as at this point the wake-optimised settings may have caused the patient to lie awake. Therefore, clinician and patient can, based on the habitual timing of the major nocturnal sleep episode, establish a personalised timed schedule, and use this to inform scheduled profile transitions. In use cases where sleep and circadian rhythms are disrupted, or where vigilance or physiological arousal are intentionally modulated with DBS, the system itself can act as a bioelectronic zeitgeber to impose a rhythm on a dysfunctional system^60^. In epilepsy, seizure probability can also show a strong daily rhythm^46–48^, and prospective stimulation adjustments might prevent the seizure occurrence rather than responding to a breakthrough.

On the other hand, a 24-hour scheduler may not capture all clinically relevant predictable variations relevant to therapy. In epilepsy, for instance, much slower infradian rhythms in seizure occurrence have been described^82^, which could impact seizure forecasting, optimal therapy level and classifier biases. Implementing infradian updates is not a fundamental limitation and can be implemented with additional memory allocation in the device scheduler. Faster rhythms include ultradian fluctuations in daytime arousal that depend on variables like sleep pressure, feeding and exercise. While the Dy2c scheduler can act with a granularity of 30 min, any stochastic or behaviour-driven ultradian rhythms could make these harder to address with the clock-based scheduler and warrant the inclusion of the reactive algorithms. Future systems with more processing resources could maintain an internal model of these variables to implement more advanced model predictions regarding therapy need and adjust algorithms accordingly (**Figure 1d**).

### 3.5 Further clinical applications

It should be noted that the relevance of time-of-day or sleep-aware brain stimulation is not restricted to our example cases of PD, epilepsy or executive attention, nor even to only nervous system disorders. Of note, sleep and circadian rhythm disruptions play an important role in psychiatric conditions^57,83–86^. DBS is now an emerging therapy in in major depressive disorder^87–89^ and may act to normalise sleep patterns^90^. In obsessive compulsive disorder, the diurnal periodicity of neural activity biomarkers has recently been directly linked to DBS effectiveness in suppressing clinical features of the disease^91^. In chronic pain, diurnal rhythms in severity are common^92–96^ and may be leveraged to provide therapy according to patient chronotype. Circadian disruption is also suggested to play a role in symptoms of urinary incontinence^97,98^, an emerging case for treatment with peripheral nerve stimulation^99^. Stimulation of the sacral, tibial, or pudendal nerves could be scheduled to follow diurnal bladder filling patterns to maximise efficacy and reduce unnecessary stimulation during sleep^100^. In a broader class of conditions, such as rheumatoid arthritis or Crohn’s disease, modulation of the inflammatory reflex via vagus nerve stimulation could be aligned with circadian peaks in the levels of pro-inflammatory cytokines, potentially optimising symptomatic control^101,102^. For cardiac pacemakers, time-based modulation of the control algorithm might align heart rate modulation with circadian autonomic tone, reducing pacing burden during sleep and enhancing physiological pacing during activity phases^103^. Finally in diabetes management, bioelectronic systems interfaced with glucose sensors and modulating autonomic pathways for insulin control could anticipate and respond more optimally to predictable daily glucose fluctuations, especially around meals or early-morning hormonal surges^104^. The inherent time-dependence of physiological systems and disorders^105^ motivates our system design for exploring new, flexible bioelectronic treatments.

### 3.6 Future outlook and conclusion

Our demonstrated ability to adapt closed-loop therapy algorithms according to time of day raises the question of what therapy settings are optimal across the diurnal cycle or for certain brain states. For example, optimal nighttime closed-loop protocols are largely unknown and may be patient specific. Even in clinically established applications such as Parkinson’s disease therapy this is currently an under-explored area, due to the limitations of available devices^34^. A device with accurate embedded real-time clock, combined with a data logger, could provide invaluable insights into intrinsic rhythms over a patient’s lifetime.

The Dy2c can serve as a ‘rheostatic research platform’ to investigate optimal parameters of time-scheduled therapy under chronic conditions, where diurnal rhythms become significant determinants of therapy effectiveness. The technical advancements demonstrated in this study provide a first step to integrating circadian phase and other time-dependent considerations with physiology-based closed-loop neuromodulation therapy on clinical-grade implantable neuromodulation systems. These results pave the way to time-dependent therapy for patients with neurological disorders whose therapy needs fluctuate throughout the day and night and brings new opportunities for investigating the brain mechanisms that regulate arousal and sleep. Devices that incorporate more physiologically-inspired algorithms may ultimately enhance therapy by adopting core control principles from nature – where evolution has optimised the balance between reactive responses and predictive adjustments for more optimal outcomes.

## 4. Methods

### 4.1 Bench testing setup

To evaluate the Dy2c system’s sensing and stimulation capabilities under controlled, physiologically relevant conditions, we used the NeuroTest platform – a custom benchtop test system developed to support the validation of adaptive sensing and stimulation systems^68,69,71^. The platform consists of an electrophoresis tank (Bio-Rad Laboratories Inc., Hercules, CA, US) filled with isotonic saline (0.9% (w/v)) and a custom hardware interface board. The saline tank serves as a model of the tissue-electrode interface, enabling realistic emulation of stimulation artefacts and their effect on sensing. During testing, the implantable pulse generator was placed centrally in the tank, with its leads positioned laterally to reflect typical clinical configurations. A reference electrode connected to the hardware interface was positioned near the device leads in the saline to record signals under conditions that approximate physiological interactions.

The NeuroTest board integrates two subsystems: a precision arbitrary current waveform generator (NeuroTest-DAC) and a high-resolution, two-channel data acquisition unit (NeuroTest-DAQ). The waveform generator supports both real-time signal generation and pre-recorded biopotential signal replay, while the acquisition unit features a 24-bit analogue-to-digital converter with a noise floor of 1 μV RMS at 1 kHz and supports sampling rates up to 32 kHz. The subsystems are galvanically isolated to reduce electrical noise, interference, and ground loop artefacts, ensuring stable and accurate characterisation. The full NeuroTest design is detailed in a recent paper^69^.

### 4.2 DyNeuMo-2c loop recorder, closed-loop operation and classifier configuration

#### 4.2.1 Loop recorder and data logging

The Dy2c’s internal memory (128 kB) can be programmed to record and stimulate in 30 min blocks over a 24-h circadian schedule. One recording and one stimulation profile can be designated for each 30 min interval. The memory buffer can hold up to 100 s worth of data sampled at 630 Hz with a signal resolution of ∼1 μV. The signal can be recorded broadband or be restricted to the power in a selected band of frequencies (power-in-band). The buffer sample storage period can go from 1.587 ms (typically for raw signals) to 1.625 s without any gaps for power-in-band signal. The buffer can store 30 hours’ worth of power-in-band signal with one full battery charge. If interruptions in the stored signal - due to battery charging - are tolerated, the buffer can record at lower rates without looping for longer time. For instance, the power-in-band signal can be recorded at a rate of 1 sample every 13 min for more than a year and 8 months. The buffer is circular, meaning that it rewrites after filling up, hence its name *loop recorder*.

#### 4.2.2 Closed-loop operation of the Dy2c

The power-in-band signal can be used to trigger closed-loop DBS when the signal crosses a preset threshold. The Dy2c will log the total number of threshold crossings during each 30 min interval, but not the timing of each crossing. Alternatively, the Dy2c can be configured to record two power-in-band signals with no stimulation or wirelessly stream a single broadband signal (up to 1 kHz) to an external PC using the handheld patient controller (Picon) and the PicoPC software. The Dy2c also has an onboard accelerometer/motion sensor that can be programmed in the scheduler to trigger changes in stimulation parameters.

#### 4.2.3 Classifier configuration with OxCAT tool

To streamline configuration of the embedded signal processing chain and classifier on the Dy2c system the OxCAT software workflow was developed^68^. Provided with labelled biopotential data, the goal of OxCAT is to dynamically select candidate filter and classifier parameters for the device embedded signal chain that maximise the classifier’s performance on the labelled input data. The OxCAT uses Bayesian optimisation to select candidate filter parameters that are simulated on the input data with the goal of maximising the Dy2c classifier precision-recall area under curve (AUC) score on the labelled dataset. If the generated classifier’s performance is sufficient, the OxCAT also enables the associated filter and classifier parameters to be exported as a configuration file for upload on to the device to enable adaptive therapy delivery based on power-in-band variations in the recorded biopotential signal.

### 4.3 Physiological models for bench testing

#### 4.3.1 Parkinson’s disease model

A Parkinsonian LFP signal was emulated using the NeuroTest platform as pseudo-random bursts of a 20 Hz oscillation, with length varying between 0.25–3 s. The frequency and amplitude of these 20 Hz ‘beta-bursts’ were modulated according to the diurnal profile observed in a real patient (**Figure 3b–d**) as previously reported^29^. Periods in the patient-recorded data with the highest power peaks resulted in more frequent and higher-amplitude 20 Hz beta bursts, sufficient to trigger the closed-loop algorithm threshold.

#### 4.3.2 Epilepsy model

Seizure data previously recorded as part of clinical practice by King’s College Hospital (London, UK) from a patient with generalised epilepsy were used for the bench testing. The patient had been implanted with externalised deep brain stimulation electrodes targeting the CMT and consented to the use of data for research purposes. The original dataset included synchronised scalp EEG and thalamic LFP recordings. Only the EEG data were used by a trained clinician to annotate periods of electrographic seizure activity, defined as epileptiform discharges lasting longer than 3 s.

A total of 48 hours of data were analysed, identifying 208 seizure events. The LFP data were pre-processed by resampling the signal to a rate of 630 Hz. To reduce computation time for bench testing, a one-hour segment containing 19 labelled seizure events was selected. From this segment, data were sliced to include each seizure event along with 10 s of signal before and after each event. These segments were then recombined into a shorter dataset. This LFP dataset was subsequently used as input to the NeuroTest platform to evaluate the performance of the Dy2c in detecting and responding to seizure activity. For illustrative purposes, 10 seconds of seizure activity and 10 seconds of pre-seizure activity (**Figure 4c**) and 10 minutes of daytime baseline CMT LFP and 10 minutes of night-time sleep CMT LFP activity (**Figure 4d**) were selected from the main data set.

### 4.4 *In vivo* validation in a healthy non-human primate

#### 4.4.1 Work involving animals

All animal work was carried out in strict accordance with the NIH Guidelines for Use of Animals in Research, under a protocol reviewed by the Weill Cornell Medical College Institutional Animal Care and Use Committee. The protocol was also approved by the Committee for Animal Care and Ethical Review at the University of Oxford. Animals were cared for by the Research Animal Resource Center at Weill Cornell Medicine.

#### 4.4.2 Cephalic implant and imaging

One adult male *Macaca mulatta* (15 kg) was implanted with a custom neuromodulation platform (Gray Matter Research LLC, Bozeman, MT, USA) to chronically record large-scale electrophysiological signals from cortical and subcortical areas and to deliver electrical stimulation to the central thalamus. The implant consists of a 4 mm thick titanium baseplate (**Supplementary Figure 1a**) that tightly conforms to the surface of the skull. The baseplate has four preset craniotomy ports to access prefrontal and thalamic targets. Following osseointegration of the baseplate and behavioural training (∼9 months later) four craniotomies were performed through the baseplate ports and a multi-chamber component, machined out of polyetherketoneketone (PEKK), was mounted to the baseplate, along with a rubber gasket to seal the craniotomies. Two custom electrode arrays (AirRay; CorTec GmbH, Freiburg, Germany) each with 32 contacts (0.79 mm^2^ contact area) that were arranged in a 4×8 square grid (20×20 mm), were placed epidurally over left and right prefrontal, premotor and motor regions of cortex (**Supplementary Figure 1b**). Four segmented DBS leads (0.85 mm diameter; Heraeus Medevio, Hanau, Germany) with 12 contacts (0.37 mm^2^ contact area) in a 4×3 ring arrangement (0.5 mm height, with inter-ring spacing of 0.5 mm) were placed in the left and right thalami to target rostral and caudal regions of the central lateral and lateral medial dorsal nuclei (**Supplementary Figure 1c**). Brass screws that passed through the PEKK chamber were threaded into the baseplate to provide ground connections for the stimulation and recording devices. Four titanium bolts were screwed into the PEKK chamber component to secure a circular ‘halo’ frame that rigidly holds the animal’s head in a docking station in the behavioural testing room. An impact resistant 3D-printed plastic cap is used protect the internal ECoG and DBS connectors (Omnetics Connector Corp., Minneapolis, MN, US) and securely hold instruments (**Supplementary Figure 1d**) during free behaviour. The implantation procedure followed previously reported methods^106^.

To plan the surgical implantation of the baseplate, ECoG arrays and DBS leads, the animal was anesthetised and placed in a magnetic resonance (MR) compatible stereotaxic surgical frame (1430M, Kopf Instruments, Tujunga, CA, US) for presurgical MR (3T Siemens) and CT (Siemens PET/CT) imaging. These images were then registered to a high-resolution MRI-DTI macaque full-brain atlas with 241 segmented anatomical structures^107^. Segmentations of the cortical areas, thalamic nuclei and models of the ECoG array and DBS leads were co-registered to the animal’s MR/CT imaging using 3D Slicer^108^. Following implantation, the locations of the ECoG arrays and DBS leads were reconstructed by registering postop CT scans within the surgical planning software and a custom SCIRun application (https://scirun.org/) to develop finite element models of the DBS leads and thalamic nuclei, as detailed in prior studies^72,109^.

#### 4.4.3 Use of the Dy2c in NHP experiments

For our long-term (> 1 h) recordings, the Dy2c was programmed to record an input signal (single dipole) from upper and lower rings of contacts from the right DBS lead or from the two sets of contacts of the right ECoG array (**Supplementary Figure 1**). The signals were conditioned on board the Dy2c as follows. 1) A first-order IIR filter was used to remove channel offset. 2) A fourth-order IIR band-pass filter was used to extract signal components in either the gamma (55– 65 Hz) or theta-alpha (5–10 Hz) band. 3) A rectifier was used to extract a noisy envelope of the filtered signal, 4) A low-pass filter, implemented as an exponential moving average, was used to obtain a smooth estimate of the envelope, resulting in the power-in-band signal.

One of the goals of this study was to demonstrate the use of the Dy2c for aDBS, which requires that we show how electrical stimulation in the central thalamus produces changes in behaviour and neural activity. The Dy2c was programmed, using the 24-hour scheduler, to provide either ‘SHAM’ stimulation, where the baseline (subthreshold) stimulation level does not change, or ‘ON’ stimulation, where stimulation is ramped up from baseline to a higher level. The duration of stimulation is controlled by setting the ‘debounce’ time for the device. Stimulation in this study was restricted to a periodic high frequency square wave at 126 Hz with a 100 µs pulse width and was ramped up from baseline at 0.01 mA per pulse. The stimulation was confined to contacts 2 and 3 of the right DBS (**Supplementary Figure 2c**). During the aDBS experiments the baseline stimulation was 0.1 mA and when triggered, the amplitude of stimulation ranged from 2.5–4.0 mA. When triggered, the system employs a timing interlock called the debounce period which ensures that stimulation state remains constant for a minimum period of time, 10–120 s in this study. We also conducted bilateral stimulation during the night and 2.0–2.5 mA consistently woke the animal up in the first half of the night, and the general arousal effect gradually diminished in the latter half of the night. However, waking the animal from sleep was not the goal of this study, and so we only present data during a unilateral protocol.

#### 4.4.4 DyNeuMo-X device

To validate the DyNeuMo-2c algorithm, we relied on a secondary recording device, denoted ‘DyNeuMo-X’. The DyNeuMo-X was designed as a more flexible research-only analogue of the Picostim–DyNeuMo devices for externalised neuromodulation studies. The system consists of a main data-logger unit, with stackable extension modules that each add recording or stimulation channels. Recording modules are based on a high-resolution data converter (ADS1299; Texas Instruments, Dallas, TX, US), supporting 8 channels per card (± 187.5 mV range, 24-bit resolution, < 1 μV_RMS_ noise floor). The data-logger integrates an ARM Cortex M7 microcontroller module (Teensy 4.0; PJCR LLC, Sherwood, OR, US) for high-throughput data processing and flexible algorithm development, battery management, a high-precision real-time clock module (RV-3028-C7; Micro Crystal AG, Grechen, Switzerland), a 3-axis accelerometer with integrated motion classifier (ADXL346; Analog Devices Inc, Wilmington, MA, US), and an LED driver for untethered synchronisation with external systems. Data is stored on exFAT-formatted microSD cards (up to 128GB).

In this study, the DyNeuMo-X was configured with a single 8-channel recording module (1 kHz synchronous sampling, broad-band), without a stimulation module. The unit was fitted with a rechargeable 1400 mAh single-cell Lithium-Polymer battery, supporting up to 13 hours of data collection. The complete device was housed within a 42×42×32 mm protective case (ESD Resin; Formlabs Inc, Somerville, MA, US). The animal’s protective cap was designed to incorporate this unit, while keeping wires and connectors secure (**Supplementary Figure 1d**).

The DyNeuMo-X was synchronised with infrared video using an LED indicator (∼1 s resolution). Importantly, common-mode signals such as power line noise (60 Hz) or abrupt motion artifacts were highly rejected by the system and were not significantly observed.

The extensive broad-band recordings acquired with the DyNeuMo-X allowed for time-frequency analysis. The power spectrum of the cortical and thalamic signals was estimated using the multitaper method as implemented in the Chronux toolbox^110^.

#### 4.4.5 Biomarker validation

As the gamma-band biomarker (55–65 Hz) overlaps with mains power frequency, several control experiments were conducted to validate the neural origin of this signal. Significant effort was put into excluding power line noise in the animal’s home environment, including disconnecting all devices in the room from mains power. Additionally, we conducted simultaneous recordings with a twin model system to have an accurate reference of baseline level of noise in the environment. A second Dy2c device was connected to either a set of DBS leads or an ECoG array identical to the ones implanted. These ‘control’ electrodes were submerged in a 0.9% saline solution, and placed adjacent to the home environment, in the same room. These tests confirmed that the regularly occurring periods of enhanced gamma-band activity were of neurophysiological origin, as they were absent in the twin system.

The regularity of these prolonged gamma-band (55-65 Hz) events allowed us to take the following approach: 1) First we used the DyNeuMo-X recordings with a computer emulation of the Dy2c signal processing chain to derive a threshold for the aDBS classifier targeting the gamma-band events (**Supplementary Figure 2c**). 2) Using this filter in the Dy2c, recorded the gamma-band (55–65 Hz) signal from the central thalamus during the night (**Supplementary Figure 2f**). The signal largely mirrored the computer emulation.

#### 4.4.6 Activity monitoring

As part of the validation of the Dy2c, we needed to evaluate the animal’s behaviour in its home environment. We sought to measure the overall motion of the animal using a 3-axis motion data logger, the AX3 (Axivity Ltd, Newcastle upon Thyne, UK). The AX3 autonomously collects accelerometer signals that can be synchronised post-hoc with the Dy2c loop recorder and infrared video. The AX3 (23×32.5×7.6 mm) was secured within the upper part of the protective cap (**Supplementary Figure 1d**) using set screws and placed adjacent to the Dy2c. The internal clock of the AX3 was updated to the PC clock when programming the onset and offset times. After an experimental session, the AX3 was removed from the protective cap and the recorded accelerometer signals were downloaded to a computer via a USB cable.

Overall acceleration magnitude was calculated by combining the acceleration values from each axis of the AX3, collected at 100 Hz, with a ±8 g resolution. The acceleration values were first squared, then summed, and the square root was taken to provide total acceleration or overall motion for each time point. A Butterworth low-pass filter (fourth order) with 5 Hz cut-off frequency was applied to attenuate high-frequency noise that partially remove large spikes, produced by collisions of the protective cap within the animal’s environment, that appeared simultaneously in all three axes and exceeded the sensor range. The final motion signal was normalised to the peak motion value during each session, which usually occurred during the morning hours from 06:00–08:00 when the lights in room turned on and husbandry staff entered the room.

#### 4.4.7 Home environment setup and video monitoring

The animal’s home environment consisted of three upper and three lower interconnected sections, each with 5.3 cubic feet of space, for a total of 31.8 ft^2^. The animal cohabited this space with another juvenile male macaque. Laboratory lights cycled between daytime (06:00–18:00 lights ON) and nighttime (18:00–06:00 lights OFF) regimes. Certain experiments required confining the animal to a sub-section of the home environment for video monitoring. In these experiments a sliding door with grooming bars was used to separate the two animals, with separate access to food and water made available. Following each 12-hour isolation period the animal was able to cohabit the entire space with his roommate again. Each animal had access to toys and a mirror, and had visual, auditory and olfactory contact with the other animal throughout the duration of the study. The animal was weighed weekly to maintain healthy weight during the study.

A wide angle (170°) infrared video camera (ELP-USBFHD05MT-KL156IR; Ailipu Technology Co, Shenzhen, China) was selected to monitor a sub-section of the home environment. The placement of the camera allowed for a full view of the animal in most conditions, except one corner of the environment where only a partial view was guaranteed. The video stream was recorded (640×480 pixels, 10 Hz) using the Synapse software and an RZ2-4 signal acquisition unit (Tucker Davis Technologies, Alachua, FL, US). Timing in the Dy2c device was synchronised post hoc (1 s resolution) with the video recordings by noting the clock time of the TDT computer.

#### 4.4.8 Stimulation methods and protocols

Following implantation of the DBS leads, we reviewed each contact to determine which contacts promoted consistent arousal. For the monopolar review we used an external stimulator (STG4004; Multichannel Systems GmbH, Reutlingen, Germany) to deliver current controlled stimulation (0.25–3.0 mA, 100μs pulse width, 150 Hz, mono- and bipolar configurations) while the animal was in the laboratory’s recording booth. Stimulation amplitudes were subsequently titrated and contact selections that promoted arousal and avoided confounders, such as motor side-effects, were used in this study. Adverse effects included brief saccadic eye movements and/or fixed gaze, contralateral to the side of stimulation. This occurred in a subset of contacts in both the left and right DBS leads. Uncontrolled hand, arm, foot, lower leg and body movements, contralateral to the side of stimulation, were also observed in a subset of contacts across the leads. Most of these adverse effects co-occurred with increased arousal, as measured by immediate pupil dilation and rapid saccadic eye movements. However, a subset of electrode contacts produced consistent pupil dilation and eye opening during drowsy periods, without any noticeable adverse effects. These contacts were then explored further using investigator-initiated periods of stimulation (10–30 s) when the animal became visibly drowsy by exhibiting slow rolling eye movements, long blinks (0.5–1 s) and full eye closures (> 1 s). Bipolar stimulation within the upper three contacts (2, 3 and 4) in one of the right DBS leads resulted in consistent eye opening, pupil dilation and increased saccadic eye movements, without noticeable side effects, up to 4.0 mA. The same stimulation parameters used by the Dy2c in this study (126 Hz, 100 μs pulse width, 0.01 mA per pulse ramp, 0.1–4.0 mA) confirmed all the above effects. Of note, the ramping capabilities (0.01 mA per pulse) of the Dy2c device markedly reduced the abrupt onset of the adverse effects, compared to the STG4004 external stimulator which didn’t ramp the level of stimulation.

To synchronise the data streamed or recorded by the Dy2c as the animal performed various behaviour tasks, brief marker signal was injected using the STG4004 stimulator. The signal consisted of a 7 Hz sinusoid packet lasting 1 s, at a sub-threshold intensity of 1 mA, delivered between the upper and lower contacts of the right DBS lead. This pattern was applied before and after the vigilance task sessions.

The Dy2c device has a compliance voltage of ± 15 V and the maximum impedance for each of the eight contacts is restricted to less than 4 kΩ. While in use, the impedance of each contact is regularly and automatically checked (∼1 s) to adjust the voltage of the stimulation engine to maintain constant current output by the Dy2c. In this study, a maximum current of 4.0 mA was used. The surface area of individual ECoG and DBS contacts are relatively small (0.79 mm^2^ and 0.37 mm^2^, respectively) compared to commercial systems, resulting in higher than usual impedances, 5–40 kΩ. To comply with the Dy2c’s impedance threshold, three neighbouring ECoG contacts (**Supplementary Figure 1b**) and the three contacts for each of the four DBS rings were combined in parallel to effectively reduce the impedance to approximate commercial systems, 0.8–1.5 kΩ.

The measured impedance of the four DBS lead rings (3 annular contact segments per ring) would occasionally rise to within 1 kΩ of the 4 kΩ upper limit of the Dy2c system. To reduce these impedances, a brief 1 Hz sinusoid stimulus of 1.0mA was applied for a duration of 4 s individually to each of the 12 contacts, based on a prior study^111^. This ‘rejuvenation’ protocol consistently reduced the measured impedance for each ring of DBS contacts to 0.6–1.2 kΩ, lasting 1–2 weeks. The measured impedances of the ECoG contacts never exceeded 2.5 kΩ.

#### 4.4.9 Vigilance task and performance monitoring

The animal was trained to perform several visuomotor reaction time tasks for liquid rewards, as done in prior studies^72,112^. First the animal’s head was fixed to a Halo-system^106^ to remove the protective cap and expose the DBS and ECoG connectors. Head fixation was also done so that it was possible to track horizontal and vertical eye movements, along with pupil diameter, using an eye tracking system (Oculomatic Pro v1.9.7, Neuro-Software Developers Inc., Jacksonville, FL, US) placed just below the LCD visual display monitor. The animal was required to initiate each trial of the task by touching and holding an infrared (IR) switch (Crist Instruments Company Inc., Hagerstown, MD, USA) mounted to the behavioural chair, which provided programmed TTL pulses to a computer controlling the task. The vigilance task was designed in NIMH MonkeyLogic^113^ which controlled the visual appearance of a central greyscale Gábor-patch (3° of visual angle) that was embedded in a 1/f noise pattern on a 20” LCD screen (60 Hz refresh rate) placed 100 cm in front of the animal.

During each trial, the animal focused on an invisible window (dotted red circle in **Figure 2a**) centred on the Gábor-patch. The spatial phase of the Gábor-patch changed (every 300 ms) for a variable number of phase transitions (lasting 1–8 s) before changing orientation (from 0° to 11.5– 22.0°). The change in orientation was the ‘GO’ cue and the animal had to quickly remove its hand from the IR switch (within 1 s) to receive a liquid reward (0.1–0.4 ml). A trial was considered to have been ‘engaged’ when the IR switch was triggered, regardless of the outcome, either correct or incorrect. For example, an incorrect trial could occur if the animal touches the switch initiating the task sequence but then breaks fixation aborting the trial; or the animal might maintain fixation and remove its hand from the IR switch shortly after triggering the task sequence; or it may initiate a trial and never remove its hand in time. Although such trials differ in terms of what the animal sees and how it moves its hand, the trials all require sustained attention and stable fixation over variable delay periods and over extended periods of time, usually lasting 1–2 hours. The sessions were stopped when the animal refused to engage in the task and typically made vocalisations and/or fell asleep for several minutes. At this point, the total water dispensed was carefully measured and the animal was moved back to its home environment.

To provide an estimate of the animal’s performance throughout the vigilance task, the series of correct and incorrect trials (assigned values of 1 and 0 respectively) were used to generate a state space model^114^. This provided a smooth estimate of percentile performance (0–100%) and was used to visualise task performance as a function of trial number.

#### 4.4.10 Eye tracking

Pupillometry measurements, along with vertical and horizontal eye movements were recorded from the previously described eye tracking system. A simple blink detection algorithm was developed in Matlab (2024b; MathWorks Inc., Natick, MA, US) to quantity the duration of eye closures, as described in a prior study^72^. The pupillometry signal was normalised to a range of 0–100%, where 100% represents fully visible pupils (eyes fully open), to 0% where the eyelids were completely closed. The pupillometry signal was then thresholded at 10% to estimate the duration of eye closures.

The resulting signal was a useful marker for interpreting neural activity. Preceding full eye closures lasting longer than 1–2 s, slow rolling eye movements and partial eye lid closures are associated with markedly enhanced spectral power in the 12–18 Hz band in the ECoG signals – just prior to full eye closure – immediately followed by shift in spectral power below 12 Hz. When the eyes re-opened (see inset in **Supplementary Figure 3c**) the power in the lower frequency bands diminished. In this study, we were interested in eye closures that lasted longer than 1 s and decided to focus on the 5–10 Hz band, as such activity has been associated with ‘microsleep’ and transitions from wake to N1 sleep in prior studies^115,116^. The anteriorisation of the alpha rhythm is a well-recognised EEG signature of drowsiness both in and outside of anaesthesia^117,118^. Of note, our frequency band of interest (5–10 Hz) spans the canonical theta (4–7 Hz) and alpha (8–12 Hz) bands in EEG literature.

## Acknowledgements

Device development and bench validation were supported the following grants: Royal Academy of Engineering Chair in Emerging technologies to TD; Medical Research Council grants MC_UU_00003/3 to TD and MC_UU_00003/6 to AS; NIHR i4i NIHR205474 to TD, MT, JJvR, RP and JF; Oxford-MRC DTP Enterprise (Industrial CASE) Studentship co-sponsored by Amber Therapeutics to KL. AD was supported by an NIHR Academic Clinical Fellowship. DJD was supported by the NIHR Oxford Health Biomedical Research Centre NIHR203316 and the UK Dementia Research Institute (UKDRI-7206) through UK DRI Ltd., principally funded by the Medical Research Council, and additional funding partner Alzheimer’s Society.

The *in vivo* work with a non-human primate was solely funded by the National Institute of Neurological Disorders and Stroke grant NS111019 (JB, KP, J-WR, NS).

We thank Dr. Antonio Valentin for sharing the seizure data used for the work presented in Fig. 4.

## Data availability

Data pertaining to the bench testing work are available on reasonable request from authors TD and JJvR. CMT LFP data are available on reasonable request from Dr. Antonio Valentin, Basic and Clinical Neuroscience Department, King’s College London, London, UK. Electrophysiological and behavioural performance data from the NHP experiments are available on reasonable request from author JB.

## References

1. Harmsen, I. E. et al. Clinical trials for deep brain stimulation: Current state of affairs. Brain Stimul. 13, 378–385 (2020).

2. Lee, D. J., Lozano, C. S., Dallapiazza, R. F. & Lozano, A. M. Current and future directions of deep brain stimulation for neurological and psychiatric disorders. J. Neurosurg. 131, 333–342 (2019).

3. Lozano, A. M. et al. Deep brain stimulation: current challenges and future directions. Nat Rev Neurol 15, 148–160 (2019).

4. Cagnan, H., Denison, T., McIntyre, C. & Brown, P. Emerging technologies for improved deep brain stimulation. Nat. Biotechnol. 37, 1024–1033 (2019).

5. Kühn, A. A. & Volkmann, J. Innovations in deep brain stimulation methodology. Movement Disord 32, 11–19 (2017).

6. Krauss, J. K. et al. Technology of deep brain stimulation: current status and future directions. Nat Rev Neurol 17, 75–87 (2021).

7. Bronte-Stewart, H. M. et al. Perspective: Evolution of Control Variables and Policies for Closed-Loop Deep Brain Stimulation for Parkinson’s Disease Using Bidirectional Deep-Brain-Computer Interfaces. Front. Hum. Neurosci. 14, 353 (2020).

8. Marks, V. S., et al. Principles of Physiological Closed-Loop Controllers in Neuromodulation. arXiv (2025) doi:10.48550/arxiv.2508.11422.

9. Conant, R. C. & Ashby, W. R. Every good regulator of a system must be a model of that system. Int. J. Syst. Sci. 1, 89–97 (1970).

10. Tinkhauser, G. & Moraud, E. M. Controlling Clinical States Governed by Different Temporal Dynamics With Closed-Loop Deep Brain Stimulation: A Principled Framework. Front. Neurosci. 15, 734186 (2021).

11. Moore-Ede, M. C. Physiology of the circadian timing system: predictive versus reactive homeostasis. *Am. J. Physiol.-Regul.*, Integr. Comp. Physiol. 250, R737–R752 (1986).

12. Stevenson, T. J. On Rheostasis. 146–166 (2023) doi:10.1093/oso/9780197665572.003.0008.

13. Hastings, M. H., Maywood, E. S. & Brancaccio, M. The Mammalian Circadian Timing System and the Suprachiasmatic Nucleus as Its Pacemaker. Biology 8, 13 (2019).

14. Cajochen, C. & Schmidt, C. The Circadian Brain and Cognition. Annu. Rev. Psychol. 76, 115–141 (2025).

15. Franken, P. & Dijk, D.-J. Sleep and circadian rhythmicity as entangled processes serving homeostasis. Nat. Rev. Neurosci. 25, 43–59 (2024).

16. Meyer, N., Harvey, A. G., Lockley, S. W. & Dijk, D.-J. Circadian rhythms and disorders of the timing of sleep. Lancet 400, 1061–1078 (2022).

17. Fleming, J. E. et al. Embedding digital chronotherapy into bioelectronic medicines. Iscience 25, 104028 (2022).

18. Harmsen, I. E., Fernandes, F. W., Krauss, J. K. & Lozano, A. M. Where Are We with Deep Brain Stimulation? A Review of Scientific Publications and Ongoing Research. Stereotact. Funct. Neurosurg. 100, 184–197 (2022).

19. Zahed, H., et al. The neurophysiology of sleep in Parkinson’s disease. Movement Disord 36, 1526–1542 (2021).

20. Lyons, K. E. & Pahwa, R. Effects of bilateral subthalamic nucleus stimulation on sleep, daytime sleepiness, and early morning dystonia in patients with Parkinson disease. J Neurosurg 104, 502–505 (2006).

21. Eugster, L., Bargiotas, P., Bassetti, C. L. & Schuepbach, W. M. M. Deep brain stimulation and sleep-wake functions in Parkinson’s disease: A systematic review. Park. Relat. Disord. 32, 12–19 (2016).

22. Zuzuárregui, J. R. P. & Ostrem, J. L. The Impact of Deep Brain Stimulation on Sleep in Parkinson’s Disease: An update. J. Park.’s Dis. 10, 393–404 (2020).

23. Wadhwa, A. et al. The effects of deep brain stimulation on sleep: a systematic review and meta-analysis. Sleep Adv. 5, zpae079 (2024).

24. Oehrn, C. R. et al. Chronic adaptive deep brain stimulation versus conventional stimulation in Parkinson’s disease: a blinded randomized feasibility trial. Nat. Med. 30, 3345–3356 (2024).

25. Stanslaski, S. et al. Sensing data and methodology from the Adaptive DBS Algorithm for Personalized Therapy in Parkinson’s Disease (ADAPT-PD) clinical trial. *npj Park*.’s Dis. 10, 174 (2024).

26. Isaias, I. U. et al. Chronic adaptive versus conventional deep brain stimulation in Parkinson’s disease: a blinded randomized pilot trial. medRxiv 2025.02.20.25322374 (2025) doi:10.1101/2025.02.20.25322374.

27. Little, S. et al. Adaptive deep brain stimulation in advanced Parkinson disease. Ann Neurol 74, 449– 457 (2013).

28. Bronte-Stewart, H. M. et al. Long-Term Personalized Adaptive Deep Brain Stimulation in Parkinson Disease: A Nonrandomized Clinical Trial. JAMA Neurol. (2025) doi:10.1001/jamaneurol.2025.2781.

29. Rheede, J. J. van et al. Diurnal modulation of subthalamic beta oscillatory power in Parkinson’s disease patients during deep brain stimulation. Npj Park Dis 8, 88 (2022).

30. Baumgartner, A. J., Hirt, L., Amara, A. W., Kern, D. S. & Thompson, J. A. Diurnal fluctuations of subthalamic nucleus local field potentials follow naturalistic sleep-wake behavior in Parkinson’s disease. SLEEP zsaf005 (2025) doi:10.1093/sleep/zsaf005.

31. Cagle, J. N. et al. Chronic intracranial recordings in the globus pallidus reveal circadian rhythms in Parkinson’s disease. Nat. Commun. 15, 4602 (2024).

32. Urrestarazu, E. et al. Beta activity in the subthalamic nucleus during sleep in patients with Parkinson’s disease. Movement Disord 24, 254–260 (2009).

33. Thompson, J. A. et al. Sleep patterns in Parkinson’s disease: direct recordings from the subthalamic nucleus. J Neurology Neurosurg Psychiatry 89, 95 (2018).

34. Yin, Z. et al. Pathological pallidal beta activity in Parkinson’s disease is sustained during sleep and associated with sleep disturbance. Nat. Commun. 14, 5434 (2023).

35. Anjum, M. F. et al. Multi-night naturalistic cortico-basal recordings reveal mechanisms of NREM slow wave suppression and spontaneous awakenings in Parkinson’s disease. Nat Commun 15, 1793 (2024).

36. Zhang, G. et al. Neurophysiological features of STN LFP underlying sleep fragmentation in Parkinson’s disease. *J. Neurol., Neurosurg*. Psychiatry 95, 1112–1122 (2024).

37. Mizrahi-Kliger, A. D., Kaplan, A., Israel, Z., Deffains, M. & Bergman, H. Basal ganglia beta oscillations during sleep underlie Parkinsonian insomnia. Proc National Acad Sci 117, 17359–17368 (2020).

38. Vetkas, A. et al. Deep brain stimulation targets in epilepsy: Systematic review and meta-analysis of anterior and centromedian thalamic nuclei and hippocampus. Epilepsia 63, 513–524 (2022).

39. Gouveia, F. V., Warsi, N. M., Suresh, H., Matin, R. & Ibrahim, G. M. Neurostimulation treatments for epilepsy: Deep brain stimulation, responsive neurostimulation and vagus nerve stimulation. Neurotherapeutics 21, e00308 (2024).

40. Salanova, V. et al. Long-term efficacy and safety of thalamic stimulation for drug-resistant partial epilepsy. Neurology 84, 1017–1025 (2015).

41. Dalic, L. J. et al. DBS of Thalamic Centromedian Nucleus for Lennox–Gastaut Syndrome (ESTEL Trial). Ann. Neurol. 91, 253–267 (2022).

42. Voges, B. R. et al. Deep brain stimulation of anterior nucleus thalami disrupts sleep in epilepsy patients. Epilepsia 56, e99–e103 (2015).

43. Suresh, S. et al. Case report: Nocturnal low-frequency stimulation of the centromedian thalamic nucleus improves sleep quality and seizure control. Front. Hum. Neurosci. 18, 1392100 (2024).

44. Kremen, V. et al. A platform for brain network sensing and stimulation with quantitative behavioral tracking: Application to limbic circuit epilepsy. medRxiv 2024.02.09.24302358 (2024) doi:10.1101/2024.02.09.24302358.

45. Sheybani, L., Frauscher, B., Bernard, C. & Walker, M. C. Mechanistic insights into the interaction between epilepsy and sleep. Nat. Rev. Neurol. 1–16 (2025) doi:10.1038/s41582-025-01064-z.

46. Karoly, P. J. et al. Cycles in epilepsy. Nat Rev Neurol 17, 267–284 (2021).

47. Khan, S. et al. Circadian rhythm and epilepsy. Lancet Neurology 17, 1098–1108 (2018).

48. Daley, J. T. & DeWolfe, J. L. Sleep, Circadian Rhythms, and Epilepsy. Curr. Treat. Options Neurol. 20, 47 (2018).

49. Bergey, G. K. et al. Long-term treatment with responsive brain stimulation in adults with refractory partial seizures. Neurology 84, 810–817 (2015).

50. Skarpaas, T. L., Jarosiewicz, B. & Morrell, M. J. Brain-responsive neurostimulation for epilepsy (RNS® System). Epilepsy Res. 153, 68–70 (2019).

51. Sun, F. T. & Morrell, M. J. The RNS System: responsive cortical stimulation for the treatment of refractory partial epilepsy. Expert Rev. Méd. Devices 11, 563–572 (2014).

52. Anderson, D. N. et al. Closed-loop stimulation in periods with less epileptiform activity drives improved epilepsy outcomes. Brain 147, 521–531 (2023).

53. Tabatabaei, F. S. A. et al. Optimizing high-frequency and low-frequency deep brain stimulation parameters for drug-resistant epilepsy: Mechanisms, clinical outcomes, and future directions. Interdiscip. Neurosurg. 41, 102109 (2025).

54. Jiruska, P. et al. Update on the mechanisms and roles of high-frequency oscillations in seizures and epileptic disorders. Epilepsia 58, 1330–1339 (2017).

55. Nassan, M. & Videnovic, A. Circadian rhythms in neurodegenerative disorders. Nat. Rev. Neurol. 18, 7–24 (2022).

56. Walker, W. H., Walton, J. C., DeVries, A. C. & Nelson, R. J. Circadian rhythm disruption and mental health. Transl. Psychiatry 10, 28 (2020).

57. Wulff, K., Gatti, S., Wettstein, J. G. & Foster, R. G. Sleep and circadian rhythm disruption in psychiatric and neurodegenerative disease. Nat Rev Neurosci 11, 589–599 (2010).

58. Schiff, N. D. et al. Behavioural improvements with thalamic stimulation after severe traumatic brain injury. Nature 448, 600–603 (2007).

59. Schiff, N. D. et al. Thalamic deep brain stimulation in traumatic brain injury: a phase 1, randomized feasibility study. Nat. Med. 29, 3162–3174 (2023).

60. Deli, A. et al. Bioelectronic Zeitgebers: targeted neuromodulation to re-establish circadian rhythms. *Conf. Proc. IEEE Int. Conf. Syst., Man*, Cybern. 2023, 2301–2308 (2023).

61. Deli, A. et al. Sleep stage can be selectively modulated through targeted Deep Brain Stimulation of Brainstem Circuits. Brain Stimul. (2025) doi:10.1016/j.brs.2025.10.001.

62. Greene, A. S., Horien, C., Barson, D., Scheinost, D. & Constable, R. T. Why is everyone talking about brain state? Trends Neurosci. 46, 508–524 (2023).

63. Gilron, R. et al. Sleep-aware adaptive deep brain stimulation control: Chronic use at home with dual independent linear discriminate detectors. Front Neurosci-switz 15, 732499 (2021).

64. Smyth, C. et al. Adaptive Deep Brain Stimulation for sleep stage targeting in Parkinson’s disease. Brain Stimul. (2023) doi:10.1016/j.brs.2023.08.006.

65. Stanslaski, S. et al. A Chronically Implantable Neural Coprocessor for Investigating the Treatment of Neurological Disorders. Ieee T Biomed Circ S 12, 1230–1245 (2018).

66. Zamora, M. et al. DyNeuMo Mk-1: Design and pilot validation of an investigational motion-adaptive neurostimulator with integrated chronotherapy. Exp Neurol 351, 113977 (2022).

67. Toth, R. et al. DyNeuMo Mk-2: An Investigational Circadian-Locked Neuromodulator with Responsive Stimulation for Applied Chronobiology. 2020 Ieee Int Conf Syst Man Cybern Smc 00, 3433–3440 (2020).

68. Fleming, J. E. et al. An embedded intracranial seizure monitor for objective outcome measurements and rhythm identification. Annu. Int. Conf. IEEE Eng. Med. Biol. Soc. IEEE Eng. Med. Biol. Soc. Annu. Int. Conf. 2023, 1–6 (2023).

69. Morrow, J. K. et al. NeuroTest: a benchtop testbed for evaluating sensing-capable electrophysiology and neurostimulation systems. bioRxiv 2025.09.19.677167 (2025) doi:10.1101/2025.09.19.677167.

70. Deli, A. et al. Tailoring Human Sleep: selective alteration through Brainstem Arousal Circuit Stimulation. medRxiv 2023.01.18.23284688 (2023) doi:10.1101/2023.01.18.23284688.

71. Cho, H. et al. Development and Evaluation of a Real-Time Phase-Triggered Stimulation Algorithm for the CorTec Brain Interchange. IEEE Trans. Neural Syst. Rehabilitation Eng. 32, 3625–3635 (2024).

72. Baker, J. L. et al. Regulation of arousal and performance of a healthy non-human primate using closed-loop central thalamic deep brain stimulation. Int. IEEEEMBS Conf. Neural Eng. : Proc. Int. IEEE EMBS Conf. Neural Eng. 2023, 10123754 (2023).

73. Baker, J. L. et al. Robust modulation of arousal regulation, performance, and frontostriatal activity through central thalamic deep brain stimulation in healthy nonhuman primates. J. Neurophysiol. 116, 2383–2404 (2016).

74. Baker, J. L. et al. Regulation of arousal and performance of a healthy non-human primate using closed-loop central thalamic deep brain stimulation. Int. IEEEEMBS Conf. Neural Eng. : Proc. Int. IEEE EMBS Conf. Neural Eng. 2023, 10123754 (2023).

75. Janson, A. P. et al. Selective activation of central thalamic fiber pathway facilitates behavioral performance in healthy non-human primates. Sci. Rep. 11, 23054 (2021).

76. Gonzalez, C., Jiang, X., Gonzalez-Martinez, J. & Halgren, E. Human Spindle Variability. J Neurosci 42, 4517–4537 (2022).

77. Purcell, S. M. et al. Characterizing sleep spindles in 11,630 individuals from the National Sleep Research Resource. Nat Commun 8, 15930 (2017).

78. Andrillon, T. et al. Sleep Spindles in Humans: Insights from Intracranial EEG and Unit Recordings. J Neurosci 31, 17821–17834 (2011).

79. Goyal, A. et al. The development of an implantable deep brain stimulation device with simultaneous chronic electrophysiological recording and stimulation in humans. Biosens. Bioelectron. 176, 112888 (2021).

80. Voges, B. R. Bi-level VNS therapy with different therapy modes at night and daytime improves seizures and quality of life in a patient with drug-resistant epilepsy. Epilepsy Behav. Rep. 24, 100633 (2023).

81. Berry, R. B. et al. The AASM manual for the scoring of sleep and associated events: Rules, terminology, and technical specification. Version 2.6. (2020).

82. Baud, M. O. et al. Multi-day rhythms modulate seizure risk in epilepsy. Nat. Commun. 9, 88 (2018).

83. Plante, D. T. The Evolving Nexus of Sleep and Depression. Am J Psychiat 178, 896–902 (2021).

84. Germain, A. & Kupfer, D. J. Circadian rhythm disturbances in depression. Hum. Psychopharmacol.: Clin. Exp. 23, 571–585 (2008).

85. Jagannath, A., Peirson, S. N. & Foster, R. G. Sleep and circadian rhythm disruption in neuropsychiatric illness. Curr. Opin. Neurobiol. 23, 888–894 (2013).

86. Mirchandaney, R., Asarnow, L. D. & Kaplan, K. A. Recent advances in sleep and depression. Curr Opin Psychiatr 36, 34–40 (2023).

87. Mayberg, H. S. et al. Deep Brain Stimulation for Treatment-Resistant Depression. Neuron 45, 651– 660 (2005).

88. Holtzheimer, P. E. et al. Subcallosal Cingulate Deep Brain Stimulation for Treatment-Resistant Unipolar and Bipolar Depression. Arch Gen Psychiat 69, 150–158 (2012).

89. Lozano, A. M. et al. Subcallosal Cingulate Gyrus Deep Brain Stimulation for Treatment-Resistant Depression. Biol Psychiat 64, 461–467 (2008).

90. Rheede, J. J. van et al. Cortical signatures of sleep are altered following effective deep brain stimulation for depression. Transl. Psychiatry 14, 103 (2024).

91. Provenza, N. R. et al. Disruption of neural periodicity predicts clinical response after deep brain stimulation for obsessive-compulsive disorder. Nat. Med. 30, 3004–3014 (2024).

92. Hu, S., Gilron, I., Singh, M. & Bhatia, A. A Scoping Review of the Diurnal Variation in the Intensity of Neuropathic Pain. Pain Med. 23, 991–1005 (2021).

93. Gilron, I., Bailey, J. M. & Vandenkerkhof, E. G. Chronobiological Characteristics of Neuropathic Pain: Clinical Predictors of Diurnal Pain Rhythmicity. Clin. J. Pain 29, 755–759 (2013).

94. Mun, C. J. et al. Circadian Rhythm and Pain: a Review of Current Research and Future Implications. Curr. Sleep Med. Rep. 8, 114–123 (2022).

95. Warfield, A. E., Prather, J. F. & Todd, W. D. Systems and Circuits Linking Chronic Pain and Circadian Rhythms. Front. Neurosci. 15, 705173 (2021).

96. Burish, M. J., Chen, Z. & Yoo, S. Emerging relevance of circadian rhythms in headaches and neuropathic pain. Acta Physiol. 225, e13161 (2019).

97. Yuan, Y. et al. Sleep Patterns and Risk of New-Onset Stress Urinary Incontinence: The UK Biobank Prospective Cohort Study. Neurourol. Urodyn. (2025) doi:10.1002/nau.70134.

98. Song, Q.-X. et al. Disruption of circadian rhythm as a potential pathogenesis of nocturia. Nat. Rev. Urol. 22, 276–293 (2025).

99. Herroelen, S. et al. Pudendal nerve stimulation for treatment of lower urinary tract symptoms: A systematic review of safety, technical feasibility and clinical efficacy. Continence 11, 101685 (2024).

100. Thoma, C. The bladder sets its own clock. Nat. Rev. Urol. 11, 544–544 (2014).

101. Fagiani, F. et al. Molecular regulations of circadian rhythm and implications for physiology and diseases. Signal Transduct. Target. Ther. 7, 41 (2022).

102. Zeng, Y., Guo, Z., Wu, M., Chen, F. & Chen, L. Circadian rhythm regulates the function of immune cells and participates in the development of tumors. Cell Death Discov. 10, 199 (2024).

103. Lee, M. T. & Baker, R. Circadian Rate Variation in Rate-Adaptive Pacing Systems. Pacing Clin. Electrophysiol. 13, 1797–1801 (1990).

104. Cauter, E. V., Polonsky, K. S. & Scheen, A. J. Roles of Circadian Rhythmicity and Sleep in Human Glucose Regulation*. Endocr. Rev. 18, 716–738 (1997).

105. Ruan, W., Yuan, X. & Eltzschig, H. K. Circadian rhythm as a therapeutic target. Nat. Rev. Drug Discov. 20, 287–307 (2021).

106. Choi, J. et al. Aseptic, semi-sealed cranial chamber implants for chronic multi-channel neurochemical and electrophysiological neural recording in nonhuman primates. J. Neurosci. Methods 420, 110467 (2025).

107. Calabrese, E. et al. A diffusion tensor MRI atlas of the postmortem rhesus macaque brain. NeuroImage 117, 408–416 (2015).

108. Fedorov, A. et al. 3D Slicer as an image computing platform for the Quantitative Imaging Network. Magn. Reson. Imaging 30, 1323–1341 (2012).

109. Janson, A. P. et al. Selective activation of central thalamic fiber pathway facilitates behavioral performance in healthy non-human primates. Sci. Rep. 11, 23054 (2021).

110. Bokil, H., Andrews, P., Kulkarni, J. E., Mehta, S. & Mitra, P. P. Chronux: A platform for analyzing neural signals. J. Neurosci. Methods 192, 146–151 (2010).

111. O’Sullivan, K. P., Orazem, M. E., Otto, K. J., Butson, C. R. & Baker, J. L. Electrical rejuvenation of chronically implanted macroelectrodes in nonhuman primates. J. Neural Eng. 21, 036056 (2024).

112. Baker, J. L. et al. Robust modulation of arousal regulation, performance, and frontostriatal activity through central thalamic deep brain stimulation in healthy nonhuman primates. J. Neurophysiol. 116, 2383–2404 (2016).

113. Hwang, J., Mitz, A. R. & Murray, E. A. NIMH MonkeyLogic: Behavioral control and data acquisition in MATLAB. J. Neurosci. Methods 323, 13–21 (2019).

114. Smith, A. C. et al. A Bayesian statistical analysis of behavioral facilitation associated with deep brain stimulation. J. Neurosci. Methods 183, 267–276 (2009).

115. Hertig-Godeschalk, A. et al. Microsleep episodes in the borderland between wakefulness and sleep. SLEEP 43, zsz163 (2019).

116. Harrison, Y. & Horne, J. A. Occurrence of ‘microsleeps’ during daytime sleep onset in normal subjects. Electroencephalogr. Clin. Neurophysiol. 98, 411–416 (1996).

117. Hasan, J. & Broughton, R. Sleep onset: Normal and abnormal processes. in Sleep onset: Normal and abnormal processes (eds Ogilvie, R. D. & Harsh, J. R.) 219–235 (American Psychological Association, 1994). doi:10.1037/10166-013.

118. Feshchenko, V. A., Veselis, R. A. & Reinsel, R. A. Propofol-Induced Alpha Rhythm. Neuropsychobiology 50, 257–266 (2004).

