## Supplementary Figures for "A clinical grade neurostimulation implant for hierarchical control of physiological activity"

**Supp. Fig 1**

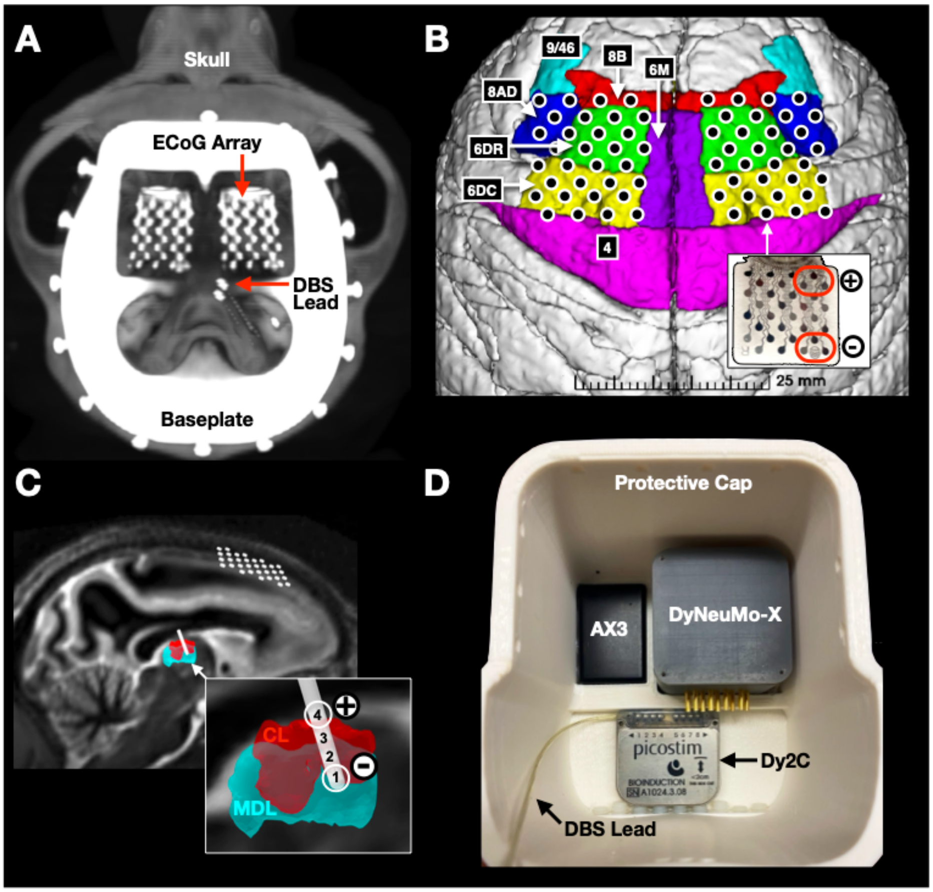


(A) A post-implant CT scan confirmed the location of the left and right ECoG arrays, and the two DBS leads place in the right thalamus. Two additional DBS leads (*not shown*) were placed in the left thalamus in a subsequent procedure. The titanium baseplate and bone screws secure the neuromodulation platform to the animal's skull.

(B) An image from the 3D slicer software used to reconstruct the individual contact locations of the left and right ECoG arrays, relative to the underlying cortical areas. A picture of the ECoG array is shown in the inset on the bottom right and the red ovals highlight the two sets of contacts, three over 8AD and three over 6DC, that were used to record prefrontal activity in this study.

(C) A sagittal image from the SCIRun software used to help plan and then reconstruct the location the DBS leads and ECoG arrays in the animal. The contacts of the right rostral DBS lead are shown relative to the targeted thalamic nuclei, the lateral medial dorsal (MDL), and central lateral (CL). The inset on the bottom right shows the two contacts, 1 and 4, that were used to record LFP activity and contacts 2 and 3 were used for stimulation.

(D) A 3D printed protective cap was designed to securely hold the AX3, DyNeuMo-X and Dy2c devices and to enclose the connections to the ECoG array and DBS lead during the home environment experiments. The view here is of the underside of the protective cap.

### Supp. Fig 2

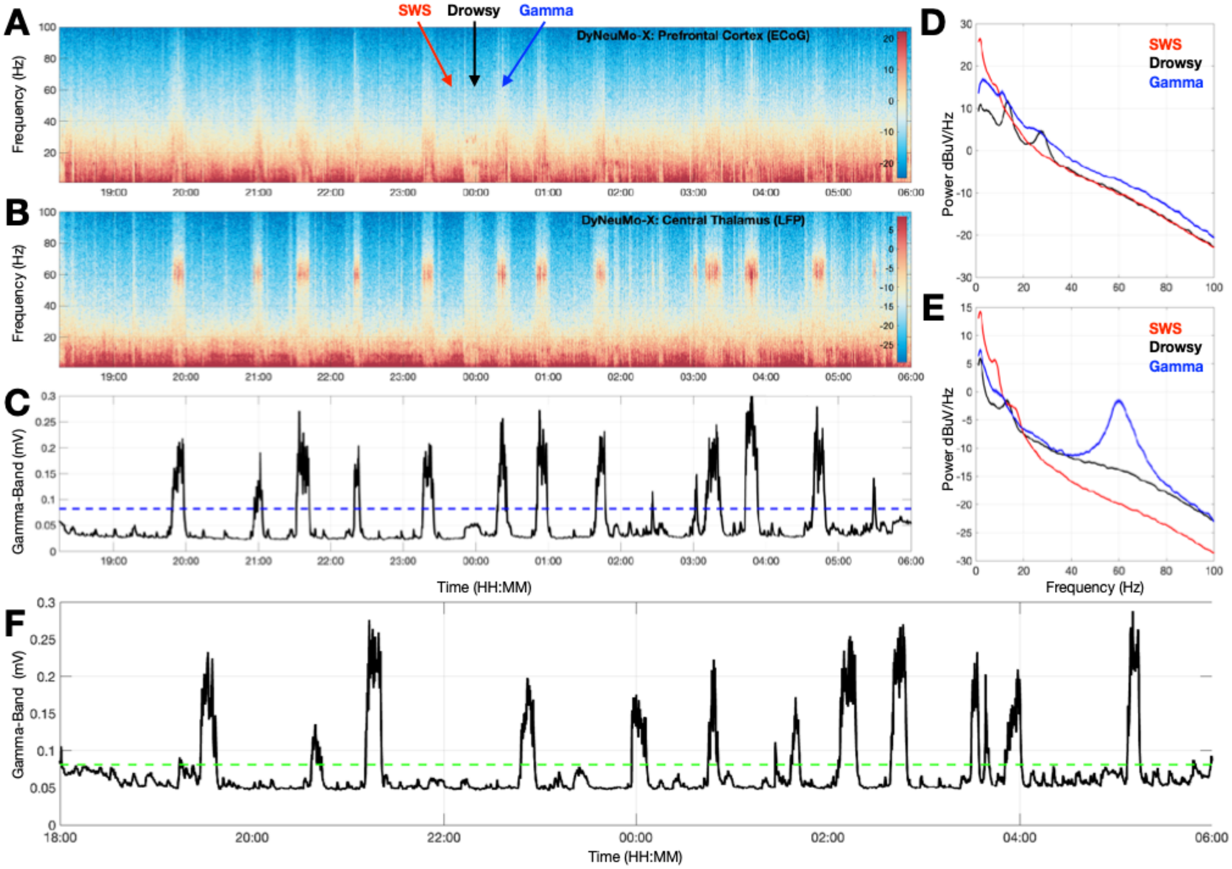


(A) A spectrogram of prefrontal ECoG activity recorder over right cortical areas 8AD and 6DC while the animal slept in its home environment during the lights OFF period from 18:00-06:00. Three 10min periods labeled ‘Gamma’, ’SWS’, and ‘Drowsy’ are marked with colored dashed lines.

(B) Same as A, but for the LFP activity recorded within the right central thalamus concurrently with the activity in A.

(C) The output of a software emulation of the Dy2c system’s power-in-band signal processing chain using a gamma-band (55-65Hz) filter for the thalamic LFP signal shown in B. The dotted green line at 0.08mV illustrates a voltage level where closed-loop CT-DBS would be triggered by the Dy2c system during *in vivo* use (see F, below).

(D) Average power spectral density plots, with 95% CI indicated by the width of the data curves, of the prefrontal cortical ECoG activity recorded during the three 10min periods marked in A.

(E) Same as D, but for LFP activity recorded currently within the right central thalamus.

(F) The animal was confined to a region of its home environment (for video monitoring), while the Dy2c recorded gamma-band (55-65Hz) activity from the right DBS lead (between contacts 1 and 4) while delivering continuous subthreshold stimulation at 0.1mA, between contacts 2 and 3, throughout the night (18:00-06:00).

### Supp. Fig 3

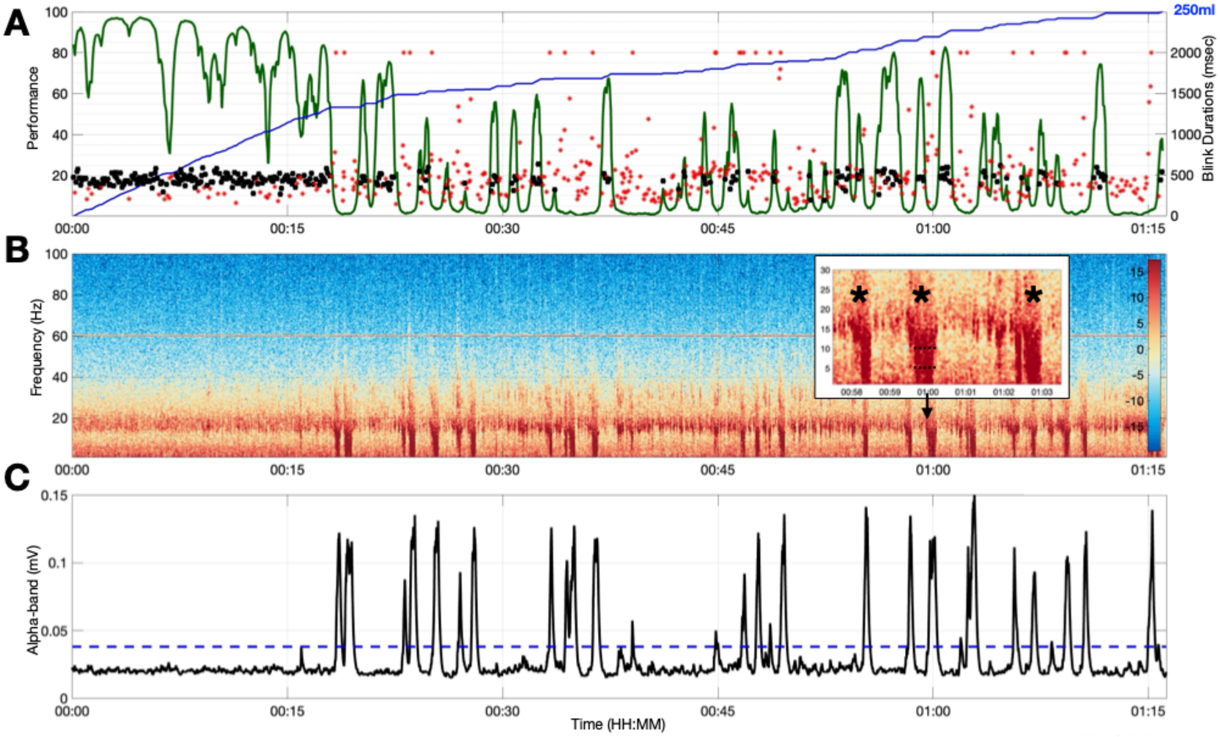


(A) The animal's performance on the vigilance task throughout one session. The green curve represents a smooth estimate of performance based on a state space model (REF). Pupillometry was used to identify the time and duration of eye blinks. Blinks that occurred during correct trials are indicated by black dots and during incorrect trials by red dots. Blinks and eye closures longer than 2sec are indicated by a ceiling of 2.0sec. The blue trace represents the accumulating liquid reward across the session (total for this day is 250ml).

(B) A spectrogram of prefrontal cortical ECoG activity recorder over right areas 8AD and 6DC was streamed by the Dy2c system while the animal performed the vigilance task. During eye closures, a significant increase in the power below 15Hz is observed, as shown in inset above the 01.00 time for three example eye closures. The 60Hz power line noise was due to the proximity of the USB powered Picon controller to the Dy2c device.

(C) The output of a software emulation of the Dy2c system's power-in-band signal processing chain using an alpha-band (5-10Hz) filter for the cortical ECoG signal shown in B. The dotted green line at 0.08mV illustrates a level where closed-loop CT-DBS would be triggered by the Dy2c system during *in vivo* use.

### Supp. Fig 4

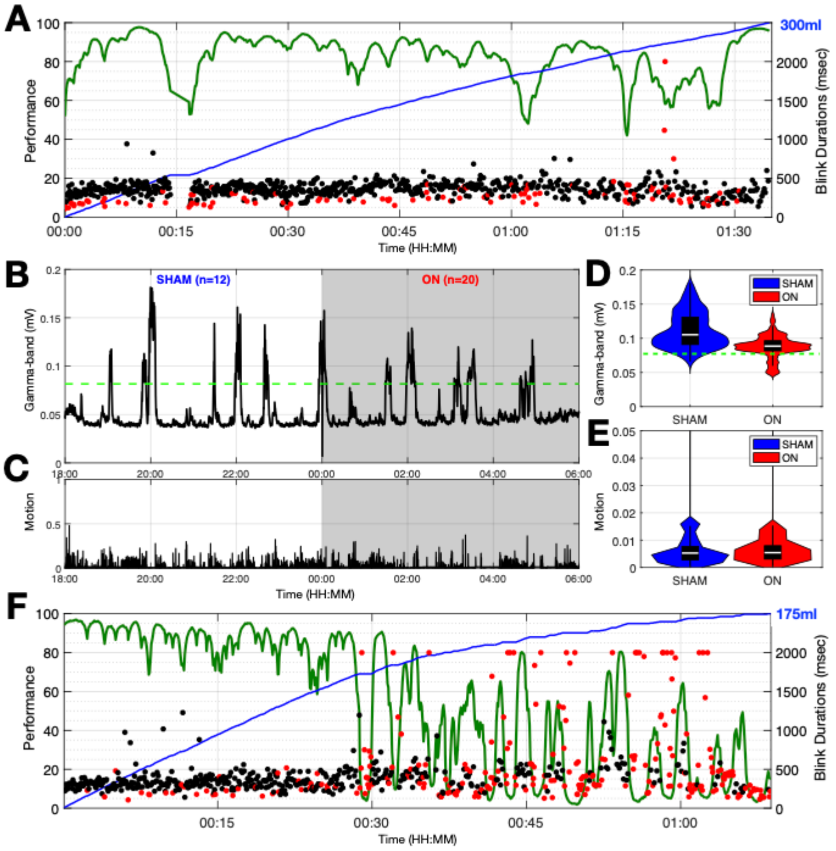


(A) The animal's performance of the vigilance task prior to a nighttime session where closed-loop CT-DBS was used to disrupt its sleep. The green curve represents a smooth estimate of performance. Pupillometry was used to identify the time and duration of eye blinks. Blinks that occurred during correct trials are indicated by black dots and during incorrect trials by red dots. Blinks and eye closures longer than 2sec are set to ceiling value of 2.0sec. The blue trace represents the accumulating liquid reward across the trial (total reward for the day's session was 300ml).

(B) The animal was confined to a region of its home environment (for video monitoring), while a closed-loop CT-DBS protocol was used by the Dy2c to continuously record gamma-band (55-65Hz) activity from the right DBS lead (between contacts 1 and 4) throughout the night. The dotted green line at 0.08mV represents the mV level in the gamma band signal used to trigger stimulation using the right DBS lead (contacts 2 and 3) that was scheduled (00:00-06:00) by the Dy2c to be ON (ramped from 0.1 – 3.0mA) or SHAM (18:00-00:00). The number of threshold crossings that occurred during the SHAM and ON conditions was logged by the device and are indicated for each 6-hour block: n = 12 for SHAM, n = 20 for ON

(C) Overall motion during the nighttime hours was derived from the 3-axis accelerometer (AX3) that was synchronized to the Dy2c loop recorder. The values here were normalized to the peak motion recorded, which usually occurred during daytime hours.

(D) Violin plots of the gamma-band (55-65Hz) signal shown in B, restricted to when closed-loop CT-DBS was triggered during the ON (00:00-06:00) and SHAM (18:00-00:00) stimulation periods. The width of the violin plots represents the probability density of the values (mV) on the vertical axis. The black box plot shows 25$^{th}$ and 75$^{th}$ percentile for each distribution, and the white line is the median value.

(E) Violin plots of the motion values during the same periods in E.

(F) The animal's performance of the vigilance task the day following the 00:00-06:00 closed loop DBS experiment shown in B.
